# A novel SATB1 protein isoform with different biophysical properties

**DOI:** 10.1101/2021.08.11.455932

**Authors:** Tomas Zelenka, Dionysios-Alexandros Papamatheakis, Petros Tzerpos, Giorgos Panagopoulos, Konstantinos C. Tsolis, Vassilis M. Papadakis, Dimitris Mariatos Metaxas, George Papadogkonas, Eleftherios Mores, Manouela Kapsetaki, Joseph Papamatheakis, David Stanek, Charalampos Spilianakis

## Abstract

Intra-thymic T cell development is coordinated by the regulatory actions of SATB1 genome organizer. In this report, we show that SATB1 is involved in the regulation of transcription and splicing, both of which displayed deregulation in *Satb1* knockout murine thymocytes. More importantly, we characterized a novel SATB1 protein isoform and described its distinct biophysical behavior, implicating potential functional differences compared to the commonly studied isoform. SATB1 utilized its prion-like domains to transition through liquid-like states to aggregated structures. This behavior was dependent on protein concentration as well as phosphorylation and interaction with nuclear RNA. Notably, the long SATB1 isoform was more prone to aggregate following phase separation. Thus, the tight regulation of SATB1 isoforms expression levels alongside with protein post-translational modifications, are imperative for SATB1’s mode of action in T cell development. Our data indicate that deregulation of these processes may also be linked to disorders such as cancer.

## 1 Introduction

In the heart of the adaptive immune system lies the proper spatiotemporal development of T cells which is coordinated by the mode of action of several transcription factors. T cell commitment during the double negative T cell stage is driven by BCL11B (Ikawa et al., 2010; Li et al., 2010a, 2010b). RUNX1 regulates cell proliferation following β-selection (Taniuchi et al., 2002; Egawa et al., 2007), upon which T cells start expressing CD4 and CD8 cell surface markers and transit into the CD4^+^CD8^+^ double positive (DP) stage (Carpenter and Bosselut, 2010). Survival and T cell maturation of DP cells are controlled by RORγt (He et al., 2000; Sun et al., 2000). Further lineage commitment into the CD4 and CD8 single positive T cell subsets is coordinated by additional factors including GATA3/TOX/ThPOK/TCF1/LEF1 for CD4 (Pai et al., 2003; Aliahmad and Kaye, 2008; Wang et al., 2008; Aliahmad et al., 2011; Steinke et al., 2014) and RUNX3 for CD8 T cells (Taniuchi et al., 2002; Egawa et al., 2007). However, the molecular mechanisms by which these transcription factors operate in a state/cell/tissue specific manner still remain elusive.

Distinct nuclear regions consisting of multiple protein molecules (key transcription factors, RNA Polymerase machinery), that interact and form compartments dubbed as transcription hubs or transcription factories (Rieder et al., 2012), have been proposed as a general mechanism of transcriptional regulation. These immobile membrane-less formations are thought to occupy specific regions of the nucleus where nucleic acids are reeled in and out instead of the proteins, to facilitate gene regulatory programs (Cook, 1999). The notion of 3D hubs has recently been proposed to describe the interplay of protein molecules with the 3D genome organization in the nucleus (Di Giammartino et al., 2020). The loop extrusion model and the cooperative role of cohesin and CTCF have been widely accepted as a dominant mechanism of genome organization and transcription regulation (Phillips and Corces, 2009; Rowley and Corces, 2018; Davidson and Peters, 2021). Apart from this, the liquid-liquid phase separation (LLPS) based model of genome organization and gene expression regulation has recently emerged, highlighting the strength of transcription factors and their domains in shaping the genome, largely driven by their capacity of interactions with each other and with nucleic acids (Boija et al., 2018; Sabari et al., 2018). LLPS is mostly driven by weak multivalent interactions between the protein’s intrinsically disordered regions (IDRs) and is often accompanied by interactions with RNA (Banani et al., 2017; Shin and Brangwynne, 2017). Low-complexity prion-like domains (PrLDs) and RNA binding domains are known features that affect a protein’s biophysical properties (Wang et al., 2018; Gotor et al., 2020). Glutamine-rich proteins and proteins harboring a poly-Q domain also display prion-like characteristics (Pearce and Kopito, 2018). Moreover, proteins with poly-Q domains and/or PrLDs are known to undergo LLPS (Zhang et al., 2015, 2020; Langdon et al., 2018; Peskett et al., 2018; Wang et al., 2018; Gotor et al., 2020). However, such proteins are also prone to abnormalities and their phase transitions frequently result in fibrous amyloids or other rigid aggregates, which are associated to several pathologies (Williams and Paulson, 2008; Peskett et al., 2018). Despite often having negative connotation, PrLDs most certainly play important physiological roles; from either regulating the solubility of proteins, to processes linked to LLPS, such as gene expression regulation and others (Franzmann and Alberti, 2019). Although the general transcription factors (CBP/p300, Mediator, RNA polymerase II), which have been largely studied in the context of LLPS, are necessary and vital for transcriptional initiation, each cell type needs the function of tissue specific pioneer factors that shape the 3D genome and regulate expression of specific cell lineages.

Our group and others have recently identified the important role of a transcription factor and genome organizer SATB1 in DP thymocytes (Feng et al., 2022; Zelenka et al., 2022), where it is predominantly expressed (Zelenka and Spilianakis, 2020). SATB1 was primarily identified as a matrix associating region (MAR) binding protein (Dickinson et al., 1992; Alvarez et al., 2000). Loss of SATB1 leads to developmental blockage at the DP T cell stage accompanied with ectopic activation of T cells in the periphery. At the molecular level, SATB1 was shown to mediate the enhancer-promoter communication of immune related genes, that are essential for the proper commitment and development of T cells (Feng et al., 2022; Zelenka et al., 2022) and its association or dissociation with these genes is regulated by its phosphorylation and acetylation, respectively (Kumar et al., 2006). Given its tight association with the nuclear matrix (Dickinson et al., 1992) there is a strong notion that one of its roles, apart from regulating transcription, is safeguarding of the nuclear structure and genomic integrity (Cai et al., 2003). This lead to the hypothesis that different SATB1 isoforms were responsible for these two distinct roles. Notably, its strong subnuclear pattern was retained even after the treatment of thymocytes with either high concentration of salt or DNase, highlighting once more its strong connection to the nuclear matrix and a possible role in nuclear structure maintenance (De Belle et al., 1998).

In this work, we utilized primary developing murine T cells, in which we have identified a novel full-length long SATB1 isoform and compared it to the canonical “short” SATB1 isoform. These isoforms displayed different biophysical properties, highlighting their potentially different roles. We showed that the interaction with RNA controlled SATB1’s localization in the nucleus. Moreover, SATB1’s phosphorylation in serine residue S635 regulated its phase separation and DNA binding ability. Consistent with its nuclear partitioning properties, SATB1 was involved in transcription and splicing and its mislocalization to cytoplasm resulted in protein aggregation. Moreover, we suggest that deregulated production of the two SATB1 isoforms and their altered phase transitions resulting in protein aggregation, can contribute to the severity of pathologies such as cancer.

## 2 Materials and Methods

### 2.1 Animals and isolation of thymocytes

All experiments were conducted in accordance with the Laboratory Animal Care and Ethics Committee of IMBB-FORTH. Animal work was approved by the IMBB Institutional Animal Care and Ethics Committee. All the experiments were performed on mice with C57BL/6 background. The generation and validation of *Satb1*^fl/fl^*Cd4*-Cre^+^ mouse was previously described (Zelenka et al., 2022). The animals used for the experiments were 4-8 weeks old. Primary thymocytes were resuspended by rubbing and passing the thymus through a 40 µm cell strainer (Falcon, 352340) in 1× PBS buffer. Cells were washed twice with 1× PBS, centrifuged at 500 g, at 4°C for 5 minutes, resuspended in 10 ml of 1× PBS and both steps were repeated.

### 2.2 *Satb1* isoforms cloning

Murine thymocytes total RNA was used for reverse transcription. The cDNA was amplified utilizing primers specific for mouse *Satb1*: SATB1-fwd (5΄-GCC <u>AGA TCT</u> ATG GAT CAT TTG AAC GAGGC-3΄, with integrated restriction enzyme site for BglII and SATB1-rev (5΄-GCC <u>CTG CAG</u> TCA GTC TTT CAA GTC GGCAT-3΄, with an integrated restriction enzyme site for PstI. The PCR product was cloned in a TOPO TA vector (Thermo Fisher Scientific, pCR™ II-TOPO®) and 20 clones with *Satb1* inserts were analyzed upon restriction enzyme digestion and Sanger sequencing indicating the presence of *Satb1* cDNAs with different length. BLAST search indicated that the cloned cDNAs encoded a short SATB1 isoform of 764 amino acids (UniProtKB – Q60611) and a long SATB1 isoform of 795 amino acids (UniprotKB – E9PVB7).

### 2.3 Custom SATB1 antibodies production

The first 330 amino acids of SATB1 short isoform (UniProtKB - Q60611, 1-764aa) fused to a 6x histidine tag at the N-terminus were expressed in *E. coli* and the antigen was utilized for the immunization of New Zealand rabbits and the production of SATB1 anti-sera (Davids Biotechnology). Rabbit polyclonal immunoglobulin specific to SATB1 N-terminal 330 amino acids were purified with column chromatography with the immobilized antigen used for immunization. The long SATB1-specific antisera were produced by Davids Biotechnology upon immunization of New Zealand rabbits with the GKGESRGVFLPSLLTPAPWPHAA peptide (corresponding to the extra peptide identified in the long SATB1 isoform) and subsequent purification of rabbit polyclonal peptide-specific immunoglobulins using affinity purification.

### 2.4 SATB1 long isoform detection and antibody validation

Thymocyte protein extracts were prepared from 4-6 weeks old female C57BL/6 mice. Thymocytes were incubated at room temperature for 30 minutes with rotation, in 1 ml EBC lysis buffer with high salt concentration (500 mM NaCl, 50 mM Tris pH=7.5, 5% Glycerol, 1% Nonidet P-40, 1 mM MgCl_2_, 1× Protease Inhibitors, 1 mM PMSF). 0.8-1 mg protein extract was incubated with 8 µg SATB1 long isoform antibody, overnight at 4°C, on an end-to-end-rotator. The immune-complexes were incubated with 20 µl protein A/G magnetic beads (Sigma Aldrich, 16-663) for 2 hours at 4°C on an end-to-end rotator. The flowthrough (SATB1 long isoform immunodepleted fraction) was kept and beads were washed 3 times with buffer I (300 mM NaCl, 50 mM Tris pH=7.5, 5% Glycerol, 0.05% Nonidet P-40, 1 mM PMSF), 2 times with Buffer II (400 mM NaCl, 50 mM Tris pH=7.5, 5% Glycerol, 1 mM PMSF) and boiled in SDS loading buffer for 10 minutes at 95°C. The flow-through was incubated with 8 µg SATB1 antibody recognizing epitopes on both isoforms, overnight at 4°C, on an end-to-end rotator. The immune-complexes were incubated with 20 µl protein A/G magnetic beads (Sigma Aldrich, 16-663) for 2 hours at 4°C on an end-to-end rotator. The flowthrough (SATB1 both isoforms immunodepleted fraction) was kept and beads were washed 3 times with buffer I (300 mM NaCl, 50 mM Tris pH=7.5, 5% Glycerol, 0.05% Nonidet P-40, 1 mM PMSF), 2 times with Buffer II (400 mM NaCl, 50 mM Tris pH=7.5, 5% Glycerol, 1 mM PMSF) and boiled in SDS loading buffer for 10 minutes at 95°C. The immunoprecipitated proteins were resolved with SDS PAGE on an 8% gel. Proteins were transferred to a nitrocellulose membrane at 320 mA for 1.5 hours at 4°C. The membrane was blocked with 5% BSA at room temperature for 1 hour and then washed three times with TBST, 5 minutes each. The membrane was initially incubated with an antibody against the long SATB1 isoform (1:200 dilution; Davids Biotechnology, custom-made), and upon antibody stripping it was blotted again using an antibody against both isoforms of SATB1 (1:100 dilution; Santa Cruz Biotechnology, sc-376096).

### 2.5 Transfection and Western Blot of recombinant SATB1 isoforms

HEK293T cells were transfected with either pEGFP-SATB1-long or pEGFP-SATB1-short plasmid vectors. Cells were lysed and whole cell protein extracts were prepared. Samples have undergone SDS PAGE and transferred in a nitrocellulose membrane for 1.5 hours at 340 mA. The membrane was blotted using SATB1 antibodies (custom-made anti-long in a 1:200 dilution and Santa Cruz sc-376097 in a 1:200 dilution) or anti-GFP (Minotech Biotechnology, 1:2000 dilution) and donkey anti rabbit-HRP (Jackson Laboratories, 1:2500 dilution).

### 2.6 Stranded-total-RNA sequencing

A biological triplicate was used for each genotype. Freshly isolated thymocytes from female animals were resuspended in 1 ml of TRIzol Reagent (Invitrogen, 15596026) and RNA was isolated according to manufacturer’s protocol. The aqueous phase with RNA was transferred into a tube and combined with 10 µg of Linear Acrylamide (Ambion, AM9520), 1/10 of sample volume of 3M CH3COONa (pH 5.2), 2.5 volumes of 100% Ethanol and tubes were mixed by flipping. Samples were incubated at – 80°C for 40 minutes. Samples were brought to 0°C and centrifuged at 16,000 g, at 4°C for 30 minutes. The supernatant was removed and the pellet was washed twice with 75% ethanol. The air-dried pellets were resuspended in 40 µl RNase-free water and incubated at 55°C for 15 minutes to dissolve. To remove any residual DNA contamination, RNase-free DNase Buffer was added to samples until 1× final concentration together with 20 units of DNase I (NEB, M0303L) and incubated at 37°C for 20 minutes. Samples were then additionally purified using the RNeasy Mini Kit (Qiagen, 74104) according to the manufacturer’s protocol. The quality of RNA was evaluated using the Agilent 2100 Bioanalyzer with Agilent RNA 6000 Nano Kit (Agilent Technologies, 5067-1511). Libraries were prepared using the Illumina TruSeq Stranded Total RNA kit with ribosomal depletion by Ribo-Zero Gold solution from Illumina according to the manufacturer’s protocol and sequenced on an Illumina HiSeq 4000 (2× 75 bp).

Raw reads were mapped to the mm10 mouse genome using HISAT2 (Kim et al., 2019). Only mapped, paired reads with a map quality >20 were retained. Transcripts were assembled with StringTie (Pertea et al., 2015) using an evidence-based Ensembl-Havana annotation file. Transcripts and genes were summarized using featureCounts (Liao et al., 2014) and statistically evaluated for differential expression using DESeq2 (Love et al., 2014, 2). When application required an intra-sample transcript comparison, DESeq2 values were further normalized to the gene length.

The list of exons was extracted from the mouse GENCODE gene set (version M20) (Frankish et al., 2019) using a customized script. Overlapping exons were merged and final exon-exon and intron-exon lists were generated. The lists were converted to 2 bp junctions (1 bp covering the first exon and 1 bp covering the second exon or the intron sequence). Both lists were intersected and purified to retain only paired exon-exon and intron-exon junctions from the same genes, i.e. embedded or overlapping genes and other unconventional cases were filtered out. Coverage of the junctions was assessed using the bedtools multicov -q 10 -s -split command and utilizing the HISAT2 mapped bam files from the RNA-seq analysis. The resulting exon-exon and exon-intron files from both WT and *Satb1* cKO were combined, junctions were paired and the unpaired junctions were filtered out. Junctions were split into the 5-prime and 3-prime end of the exon. Splicing efficiency was estimated for entire genes after summing up the coverage of all junctions from the same gene. Genes with overall exon-exon WT coverage (all exon-exon junctions together for all biological replicates) lower than 50 were filtered out. The splicing efficiency was determined as the intron/exon coverage ratio and its changes as a difference between *Satb1* cKO – WT intron/exon ratios. Moreover, the SATB1 binding sites’ dataset (GSE173446; Zelenka et al., 2022) was intersected with the junctions to search for a potential dependency of splicing on the SATB1 presence (**Supplementary Figure 4d**). The SATB1 binding propensity was calculated as the sum of fold change enrichments of all SATB1 peaks (for each peak summit against random Poisson distribution with local lambda, based on MACS2) (Zhang et al., 2008, 2) overlapping each gene and normalized to 1 kbp.

Alternative splicing was analyzed using the Whippet program (Sterne-Weiler et al., 2018) with --biascorrect flag and using both options: with and without the --bam flag (using the HISAT2 mapped bam files from the RNA-seq analysis, ensuring coverage of unannotated splice-sites and exons). In both cases, the mouse GENCODE gene set (version M11) (Frankish et al., 2019) was used to build the index file. The results from the whippet-delta script were filtered for results with probability >=0.9 and absolute delta psi value >=0.1. Out of 150,382 and 130,380 events it yielded 3,281 (2,015 genes) and 1,717 (1,077 genes) events that passed the filtering step for the bam-file-supplemented and without the bam-file analyses, respectively. Moreover, the analysis of back-splicing (circular RNA) revealed 639 events (519 genes) being underrepresented and 443 events (374 genes) overrepresented in the *Satb1* cKO.

### 2.7 SATB1 co-immunoprecipitation coupled to mass spectrometry

Thymocyte protein extracts were prepared from eight male 4-6 weeks old C57BL/6 mice. Thymocytes were incubated at room temperature for 30 minutes with rotation, in 1 ml lysis buffer (150 mM NaCl, 50 mM Tris pH=7.5, 5% Glycerol, 1% Nonidet P-40, 1 mM MgCl_2_, 200 U benzonase, 1× Protease Inhibitors, 1 mM PMSF). 24 mg of protein extract were precleared using 100 μl of Dynabeads® Protein G (Life Technologies, 10004D) at 4°C for 1.5 hours on an end-to-end rotator. 50 μl of beads pre-blocked with 15 μg rabbit anti-SATB1 or rabbit IgG sera (Santa Cruz Biotechnology, C2712) at 4°C for 4 hours rotating and crosslinked at room temperature for 30 minutes with tilting/rotation using cross-linker Bis(sulfosuccinimidyl) suberate BS3 (Thermo Fisher Scientific, 21580). Precleared lysates were subjected to immunoprecipitation by incubation with Dynabeads crosslinked with anti-SATB1 or IgG at 4°C for 30 minutes. Beads were washed three times with buffer I (150 mM NaCl, 50 mM Tris pH=7.5, 5% Glycerol, 0.05% Nonidet P-40, 1mM PMSF), twice with Nonidet P-40 free Buffer II (150 mM NaCl, 50 mM Tris pH=7.5, 5% Glycerol, 1 mM PMSF) and boiled at 95°C for 5 minutes.

Proteins co-Immunoprecipitated with SATB1 were resolved via SDS-PAGE in a 10% polyacrylamide gel. The gel was stained with silver nitrate compatible with mass spectrometry fixation: 50% MeOH, 12% acetic acid, 0.05% Formalin for 2 hours and washed with 35% ethanol for 20 minutes for 3 times. Sensitization followed with 0.02% Na_2_S_2_O_3_ for 2 minutes and washes performed with H_2_O for 5 minutes 3 times. Staining was performed with 0.2% AgNO_3_, 0.076% formalin for 20 minutes and washes were with H_2_O for 1 minute 2 times. Development took place with 6% Na_2_CO_3_, 0.05% formalin, 0.0004% Na_2_S_2_O_3_. Staining was stopped with 50% MeOH, 12% acetic acid for 5 minutes.

Ten pairs of protein bands (anti-SATB1, IgG) were excised from silver-stained polyacrylamide gels and cut into small pieces. Gel pieces were covered with destain solution (30 mM potassium ferricyanide, 100 mM sodium thiosulfate) and vortexed for 10 minutes in order to remove silver nitrate. For the reduction and alkylation of the cysteine residues, the gel pieces were covered with 100 µl 10 mM DTT and shaked for 45 minutes at 56°C followed by 45 minutes shaking at room temperature with 100 µl 55 mM iodoacetamide. Proteins were digested in 30 µl of diluted Trypsin solution and incubated overnight at 37°C. The next day, the supernatant was collected. For the extraction from the gel matrix of generated peptides, gel pieces were shaken for 20 minutes first in 50 μl 50% acetonitril (ACN) and finally in 50 μl 0.1% TFA/50% ACN. Peptide solutions were centrifuged by Speed Vac until dry powder remained and then analyzed by means of liquid chromatography combined with mass spectrometry analysis (nanoLC-MS/MS), as previously described (Stratigi et al., 2015).

Peptide intensities of each experiment were normalized based on the mean difference of their distribution (log2 intensity) with the reference sample (SATB1 IP-MS 2nd biological replicate). Missing values were imputed using the mean values of each peptide within a given study group (SATB1 IP-MS or the control – IgG IP-MS), plus random noise equal to the mean observed variability in the dataset. Dataset variability was calculated as the standard deviation of the difference between all the biological replicates of a given group and the group mean. Protein abundance was calculated using the sum of peptide intensities. Differentially abundant proteins were selected after comparing the t-test p-value (using log2 intensities) and the fold change for each protein. P-values were adjusted for multiple hypothesis testing error using the Benjamini-Hochberg method. Threshold for selection of differentially abundant proteins was an adjusted p-value < 0.05 and a fold change > 2×. Proteins that were uniquely identified in the SATB1 IP-MS dataset and proteins commonly identified with a significantly higher abundance in the SATB1 IP-MS replicates were considered for the downstream analysis as enriched proteins.

Protein-protein interaction data were downloaded from the STRING database (Szklarczyk et al., 2019). STRING database mappings from Ensemble protein IDs to Uniprot IDs were used to map the interaction data with the experimental results. Protein-protein interaction network was build using the NetworkX Python library (Hagberg et al., 2008). To cluster network nodes, vector representations were initially extracted for each node, using the node2vec algorithm (Grover and Leskovec, 2016) and followed by k-means clustering. GO and pathway enrichment analysis were performed using g:Profiler tool (Raudvere et al., 2019) using the default settings.

### 2.8 Immunofluorescence confocal and super-resolution microscopy experiments

Freshly isolated thymocytes were incubated for 60 minutes (producing the best results compared to 30 and 240 minutes) at 37°C in complete DMEM (GIBCO, 11995-073) medium with 10% serum with 2 mM 5-Fluorouridine (FU; Sigma Aldrich, F5130) or directly attached on poly-D-lysine-coated coverslips. For super-resolution microscopy, high precision Zeiss squared coverslips of 1.5H thickness (0.170+-0.005 mm; Marienfeld Superior, 0107032) were treated with 1 M HCl overnight, rinsed with ddH_2_O and then stored in absolute ethanol. Right before the experiment, coverslips were coated by dipping in 0.1 mg/ml poly-D-lysine solution (Sigma Aldrich, P6407). Attached cells were washed once with 1× PBS. Coverslips to be treated with 1,6-hexanediol (Sigma Aldrich, 804308) were incubated for 5 minutes at room temperature with the 1,6-hexanediol solution, carefully washed twice with 1× PBS and then fixed for 10 minutes on ice with 4% formaldehyde (Pierce, 28908) in 1× PBS. Fixed cells were permeabilized with 0.5% Triton-X in 1× PBS for 5 minutes on ice. Cells were washed three times with 1× PBS for 5 minutes each and blocked for 30 minutes at room temperature with Blocking Buffer [0.4% acetylated BSA (Ambion, AM2614) in 4× SSC] in a humidified chamber. Cells were incubated for 1.5 hours at room temperature with primary antibodies (see the table below) in Detection Buffer (0.1% acetylated BSA, 4× SSC, 0.1% Tween 20) in a humidified chamber. The excess of primary antibodies was washed away by three washes, for 5 minutes each, with Wash Buffer (4× SSC, 0.1% Tween 20). Cells were incubated for 60 minutes at room temperature with the secondary antibodies in Detection Buffer (0.1% acetylated BSA, 4× SSC, 0.1% Tween 20) in a humidified chamber. The excess was washed away by three washes, for 5 minutes each, with Wash Buffer (4× SSC, 0.1% Tween 20). The coverslips for 3D-SIM experiments were stained for 5 minutes at room temperature with DAPI in 1× PBS (0.85 μg/ml) and then washed three times in 1× PBS. The coverslips for STED experiments were directly mounted with 90% glycerol + 5% (w/v) n-propyl gallate non-hardening medium and sealed with a transparent nail polish. Two or three biological replicates were performed for each experiment.

Summary of antibodies used in super-resolution microscopy experiments:

| Factor | Primary antibody |  |  | Secondary antibody STED |  |  | Secondary antibody 3D-SIM |  |  |
| --- | --- | --- | --- | --- | --- | --- | --- | --- | --- |
|  | Vendor | Cat. No. | Dil. | Name | Species specificity | Dil. | Name | Species specificity | Dil. |
| long SATB1 isoform | Davids Biotechnology | n/a | 1:500 | Abberior Star Red or Alexa Fluor 555 | goat anti-rabbit | 1:100 or 1:500 | Alexa Fluor 488 | goat anti-rabbit or donkey anti-rabbit | 1:500 |
| all SATB1 isoforms | Davids Biotechnology | n/a | 1:500 | Abberior Star Red | goat anti-rabbit | 1:100 | Alexa Fluor 488 | goat anti-rabbit | 1:500 |
| all SATB1 isoforms | Santa Cruz Biotechnology | sc-5990 | 1:200 | — | — | — | Alexa Fluor 488 | donkey anti-goat | 1:500 |
| 5-fluorouridine | Sigma Aldrich | B2531 | 1:250 | Abberior Star 580 | goat anti-mouse | 1:100 | Alexa Fluor 568 | goat anti-mouse | 1:500 |

The STED microscopy images were acquired using an inverted microscope DMi8 with laser scanning confocal head Leica TCS SP8, equipped with STED 3X module, 100x/1.4 oil immersion objective and 660 nm continual and 775 pulse depletion lasers. This system enables super-resolution imaging with 35 nm lateral and 130 nm axial resolution. Depletion lasers were set to 60% and typically only a single representative z-stack from the center of each cell was scanned to reduce the effect of photobleaching and to maximize the lateral resolution, with line accumulation set to 8 and speed of 400 Hz. Raw images were deconvoluted using the Huygens Professional imaging software. The final images were analyzed using custom-made macros in Fiji software (Schindelin et al., 2012). Cells from the representative middle z-stack were manually selected for the analysis and the integrated density was calculated by the measure tool of Fiji. The pixel-based co-localization analysis of selected cells was performed using the Coloc2 plugin. SATB1 scans rotated by 90° served as a negative control for the co-localization with FU. The presented images have artificially increased contrast and brightness for better visualization.

The 3D-SIM experiment images were acquired using the DeltaVision OMX V4 imaging platform equipped with PlanApo N 60x/1.42 oil immersion objective, pco.edge 5.5 sCMOS cameras and 405, 445, 488, 514, 568 and 642 nm lasers. Spherical aberration was minimized using immersion oil with RI 1.516 for sample acquisition. Images were acquired over the majority of the cell volume in z-dimension with 15 raw images per plane (five phases, three angles), providing ∼120-135 nm lateral and ∼340-350 nm axial resolution for 488/568 nm lasers, respectively. Image reconstruction and deconvolution was performed by the SoftWoRx software using channel-specific OTFs with Wiener filter set to 0.01 for DAPI and 0.001 for the other channels. For data analysis, mostly a single representative z-stack from the center of the cell was manually selected to minimize any effect of artefacts. The final images were analyzed using custom-made macros in Fiji software (Schindelin et al., 2012). Cells were manually selected for the analysis and then the exact nuclear borders were drawn based on the DAPI signal. This was used to calculate the integrated density by the measure tool of Fiji as well as to perform the pixel-based co-localization analysis using the Coloc2 plugin. SATB1 scans rotated by 90° served as a negative control for the co-localization with FU. The DAPI zonation was achieved using the IsoData Classifier in combination with the machine learning Trainable Weka Segmentation plugin. The selection of the resulting zones was overlaid with SATB1 speckles, identified as points using Find Maxima function in Fiji with prominence set to 2000 or 5000 depending on the biological replicate. For each cell, the number of points in each zone was converted to its relative proportion in respect to the total number of speckles in the cell. The presented images have artificially increased contrast and brightness for better visualization.

For the co-localization immunofluorescence experiments analyzed with confocal microscopy (**Supplementary Figure 1f**) we utilized a commercially available antibody against SATB1 detecting both short and long isoforms (1:100; Santa Cruz Biotechnology, sc-376096) and our custom-made polyclonal antibody (1:500; Davids Biotechnology) raised against the extra SATB1 peptide of the long isoform, following the protocol described earlier. Coverslips were mounted with hardening medium ProLong Gold with DAPI (Invitrogen, P36935). Images were taken using the inverted microscope DMI6000 CS with laser scanning confocal head Leica TCS SP8, equipped with 63x/1.40 oil immersion objective.

### 2.9 Nuclear matrix preparation

Freshly isolated thymocytes were attached to poly-D-lysine coated coverslips. Soluble proteins were removed by extraction in cytoskeletal buffer (CSK: 10 mM Pipes pH 6.8, 300 mM sucrose, 100 mM NaCl, 3 mM MgCl_2_, 1 mM EGTA, 2 mM VRC) supplemented with 0.5% Triton X-100 for 3.5 minutes at 4°C. The first group of cells was cross-linked with 4% PFA in 1× PBS for 10 minutes at 4°C, followed by three washes in 70% Ethanol. The other groups were subjected to DNase treatment in 300 μl DNase Buffer (10 mM Tris-HCl pH 7.5, 2.5 mM MgCl_2_, 0.5 mM CaCl_2_) combined with DNase I (10 U = 5 μl; NEB, M0303L) at 37°C for 40 minutes. The last group was subjected to RNase A treatment (10 μl = 100 μg; Qiagen, 1007885) in PBS at 37°C for 40 minutes. After the enzymatic treatment, nucleic acid fragments were extracted by a 5-minute treatment on ice with CSKpremix (without Triton X-100) supplemented with 250 mM Ammonium Sulfate, followed by one 1× PBS wash and then they were fixed as previously described. Alternatively, they were further extracted with CSKpremix with 2M NaCl, followed by one 1× PBS wash and then fixed. After fixation, cells were washed twice with 1× PBS. Samples were blocked for 30 minutes with Blocking Buffer as previously described and stained with an anti-SATB1 antibody targeting all isoforms (Santa Cruz Biotechnology, sc-5990) and a donkey anti-goat Alexa Fluor 546 secondary antibody. Coverslips were mounted with hardening medium ProLong Gold with DAPI (Invitrogen, P36935). Images were taken using the inverted microscope DMI6000 CS with laser scanning confocal head Leica TCS SP8, equipped with 63x/1.40 oil immersion objective. Images were scanned at 600 Hz speed, unidirectionally with frame average set to 3. In the experiment with double DNase I + RNase A treatment, no cells were detectable using fluorescence microscopy, hence the remaining cellular shells were identified in bright field. The final images were analyzed using the Fiji software (Schindelin et al., 2012). The presented images have artificially increased contrast and brightness for better visualization.

### 2.10 Construction of CRY2-mCherry SATB1 recombinant proteins

For all CRY2 constructs, the pCMV-CRY2-mCherry vector was used (a gift from Won Do Heo; Addgene plasmid # 58368). As a positive control, the N-terminal part of FUS protein was used from the plasmid pHR-FUS_N_-mChr-CRY2WT (a gift from Clifford Brangwynne; Addgene plasmid # 101223). The N-terminal part of SATB1 was cloned into the pCMV-CRY2-mCherry vector using a single BspEI restriction enzyme site from the full-length construct. The IDR part of all *Satb1* isoforms (incl. the S635A mutated version) and also the N-terminal part of *FUS* was PCR amplified from plasmids, generating new restriction enzyme sites for XhoI and BamHI, using the following primers: for *Satb1*fwd: GTCTACTCGAGCACGAAAGGAAGAGGACCCC, rev: GATTGGATCCTTACACGGAAATTTTGGTTCGTG, for *FUS* fwd: GTCTACTCGAGCAATGGCCTCAAACGATTATACC, rev: GATTGGATCCTTATCCACGGTCCTGCTGTCCATAG. The sequence encoding the SV40 NLS signal peptide PKKKRKV was isolated from a lab’s plasmid using BspEI and SalI enzymes and inserted into CRY2-mCherry-IDR-SATB1 constructs between BspEI and XhoI sites, embedding it between mCherry and the IDR part of SATB1. Bacteria were grown in Luria-Bertani Broth at 37°C and plasmid DNA was isolated using the Nucleobond Xtra midi kit (MACHEREY-NAGEL, 740410), according to the manufacturer’s instructions. PCR reactions were performed utilizing the Phusion High-Fidelity PCR Master Mix (NEB, M0531L) to prevent any mutations. For all restriction reactions the respective NEB reagents were used according to the manufacturer’s instructions. Ligation reactions were performed using T4 DNA Ligase (Minotech, 202-2) according to the manufacturer’s instructions. Ligation products were transformed into DH5a competent cells via 42°C heat shock for 45 seconds. All constructs were sequence-verified using Sanger sequencing in Macrogen Europe.

### 2.11 Live cell microscopy

Live cell imaging was performed in a live imaging box at 37°C with 5% CO_2_, using the inverted microscope DMI6000 CS with laser scanning confocal head Leica TCS SP8, equipped with a 63x/1.40 oil immersion objective. To ensure the minimal pre-activation effect, cells were located with dim bright field illumination. In all cases presented, only a single plane in the center of the cell was scanned. CRY2 activation was achieved by 488 nm Argon laser illumination every two seconds, complemented with DPSS 561 nm laser illumination for mCherry detection. During the inactivation period, only the 561 nm laser was used to detect the mCherry signal. The scanning parameters were: PinholeAiry 1.5 AU, frame average 2 and speed to 600 Hz, scanned unidirectionally. Images were analyzed using custom-made macros in Fiji software (Schindelin et al., 2012) and data were processed in Bash and R. Briefly, images were corrected for movement using the StackReg plugin (Thévenaz et al., 1998). After basic pre-processing, droplets were identified in each cell for each time point using the Analyze Particles function of Fiji. The area of droplets was used to determine the droplet size. The number of droplets was normalized to the size of a nucleus, determined by the background mCherry signal in combination with bright field imaging. The images presented have artificially increased contrast and brightness for better visualization.

For the FRAP experiments, cells were first globally activated by 488 nm Argon laser illumination (alongside with DPSS 561 nm laser illumination for mCherry detection) every 2 s for 180 s to reach a desirable supersaturation depth. Immediately after termination of the activation phase, light-induced clusters were bleached with a spot of ∼1.5 μm in diameter. The scanning speed was set to 1,000 Hz, bidirectionally (0.54 s / scan) and every time a selected point was photobleached for 300 ms. Fluorescence recovery was monitored in a series of 180 images while maintaining identical activation conditions used to induce clustering. Bleach point mean values were background subtracted and corrected for fluorescence loss using the intensity values from the entire cell. The data were then normalized to mean pre-bleach intensity and fitted with exponential recovery curve in Fiji or in frapplot package in R.

### 2.12 Structure of SATB1 and the use of predictors

SATB1 domains used in **Figure 1a** were previously described (Zelenka and Spilianakis, 2020). The prediction of IDR regions in **Figure 4a** is based on PONDR VL3 score (http://www.pondr.com/) and it is shown only for the long SATB1 isoform. Similarly, the sequence of the long SATB1 isoform was analyzed using the hidden-Markov model algorithm PLAAC to predict prion-like domains (Lancaster et al., 2014), using background frequencies from *S. cerevisae*. For better lucidity, the background probabilities are not shown.

**Figure 1.**
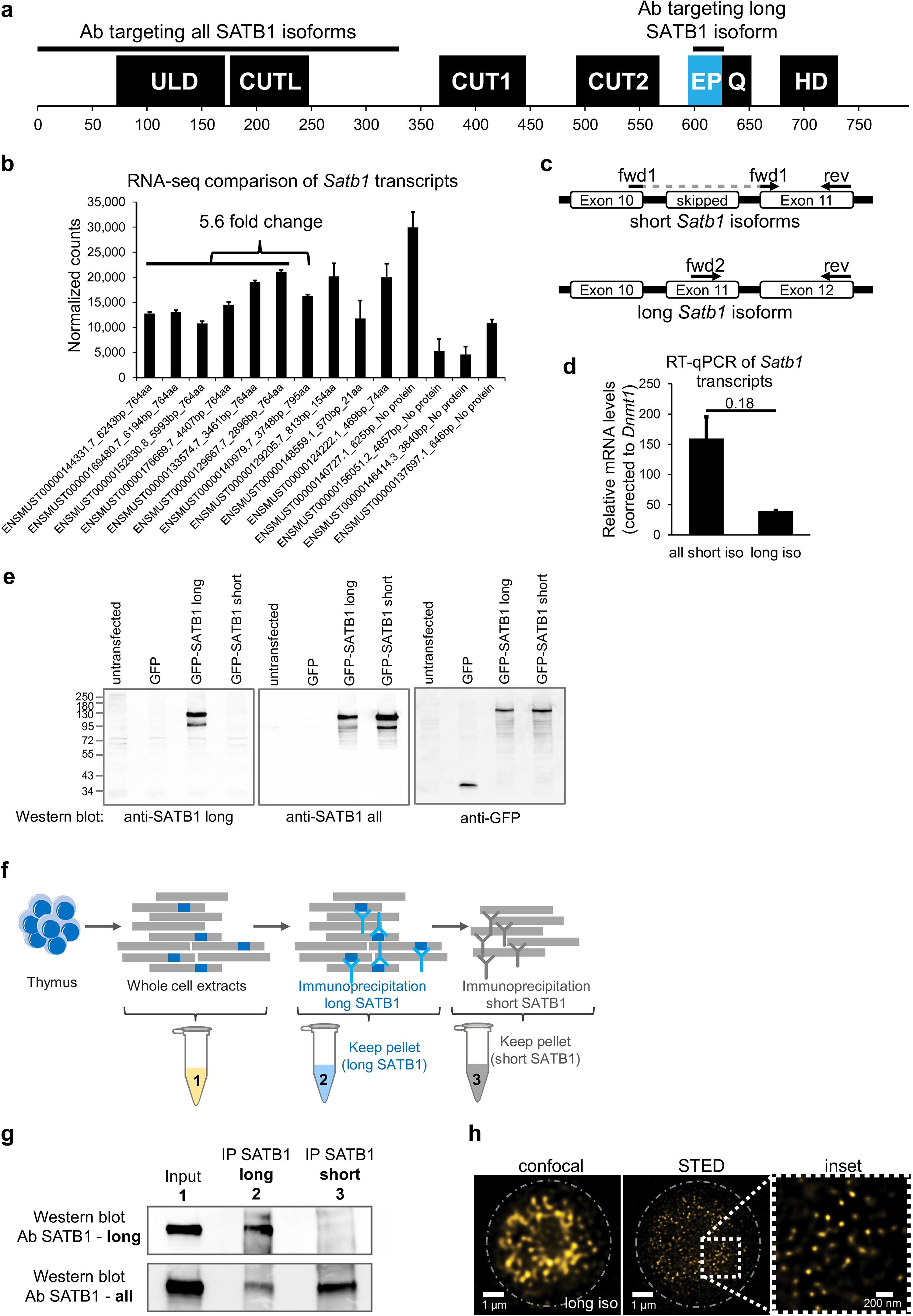
Identification of two SATB1 protein isoforms in murine thymocytes. **a**, SATB1 protein consists of the following domains and structural features: ULD – ubiquitin-like domain, CUTL – CUT-like domain, CUT1 and CUT2 domains, EP – the peptide encoded by the predicted extra exon of the long *Satb1* isoform, Q – compositional bias represented by a poly-Q domain and a stretch of prolines, HD – homeodomain. In this study, we used two custom-made antibodies (Davids Biotechnology): an antibody targeting the first 330 amino acids of all isoforms (cannot discriminate between the two protein isoforms) and the long isoform antibody specifically targeting the extra peptide of the long isoform. **b**, Comprehensive list of quantified *Satb1* isoforms based on stranded-total-RNA-seq experiment in thymocytes. Data are the mean ± s.d. **c**, Design of the confirmatory RT-qPCR experiment to quantitate relative expression of the short and long *Satb1* isoforms in murine thymocytes. **d**, RT-qPCR results for the relative mRNA levels of the two *Satb1* isoforms. Values from 3 technical and 2 biological replicates were normalized to *Dnmt1* mRNA levels. Data are the mean ± s.d. *P* values by Student’s T-test. **e**, The custom-made antibody detecting the long SATB1 protein isoform cannot detect the short SATB1 isoform. Protein extracts (80 µg) prepared from transfected HEK293 cells with plasmids expressing GFP, GFP-SATB1-long and GFP-SATB1-short. Western blotting performed with antibodies detecting either the long SATB1 isoform, all SATB1 isoforms or GFP. **f**, Scheme for the approach utilized to detect the SATB1 long isoform and validate the custom-made long isoform-specific antibody (Davids Biotechnology). Whole cell thymocyte protein extracts (sample 1) were prepared and incubated with a custom-made antibody against the long SATB1 isoform. The immunoprecipitated material (sample 2) was kept and the immunodepleted material was subjected on a second immunoprecipitation reaction utilizing an antibody detecting epitopes on both long and short SATB1 isoforms (Santa Cruz Biotechnology, sc-376096). The material from the second immunoprecipitation reaction was kept (sample 3). The second biological replicate is depicted in **Supplementary Figures 1a,b**; including the thymocyte extract immunodepleted for all SATB1 isoforms (sample 4). **g**, Western blot analysis for the samples described in **f**. The whole thymocyte protein extract (1), immunoprecipitated long SATB1 protein (2) and immunoprecipitated short SATB1 protein (3). UPPER PANEL: Western blot analysis utilizing a SATB1 antibody detecting only the long SATB1 isoform. LOWER PANEL: Western blot analysis utilizing a SATB1 antibody detecting all SATB1 isoforms. **h**, Confocal and STED microscopy images indicating the subnuclear SATB1 cage-like pattern. The super-resolution microscopy unveils that the cage-like pattern is actually composed of individual, mostly round, 40-80 nm large speckles.

### 2.13 Breast cancer data processing

Breast invasive carcinoma cohort data (BRCA) from the TCGA were obtained from the FireBrowse repository of the Broad Institute of MIT & Harvard (http://firebrowse.org). For the correlation experiment between chromatin accessibility and RNA expression, the ATAC-seq bigwig files were downloaded (Corces et al., 2018) and signal from a region encompassing the extra exon of the long *SATB1* isoform (chr3:18351000-18351700) was extracted using multiBigwigSummary from the deeptools package (Ramírez et al., 2016). The signal was correlated with summarized values of the two short isoforms (uc003cbh.2 and uc003cbj.2) and separately with the long isoform (uc003cbi.2) RNA-seq values (level 3, isoform-specific normalized RSEM dataset, version 2016_01_28). Patient TCGA-A2-A0ET-01A was removed as an outlier. The same RNA-seq values were also used for correlation with the metadata on the pathological T stage of cancer patients. Next, we analyzed the recurrent copy number alterations using the R package GAIA (Morganella et al., 2011), using the SNP6 level 3 segmented sCNA minus germline CNV dataset. The output file with tumor samples was further processed by the GenomicRanges package (Lawrence et al., 2013) and annotated by the findOverlaps function. Survival plots were constructed using the survival package in R (Therneau and Grambsch, 2000), utilizing the normalized RNA-seq data and clinical metadata from TCGA. RNA-seq data were transformed into z-scores as follows: z = [(value gene X in tumor Y) – (mean gene X in all tumors with CNV segment mean value <|0.2|)] / (standard deviation X in all tumors with CNV segment mean value <|0.2|). Patients were divided into groups with low and high estrogen receptor levels. Moreover, patients were divided based on the relative expression levels of each tested isoform into UPregulated, DOWNregulated and unchanged groups, using the z-score threshold 1.65 (∼ *P* = 0.0495). Survival plots for individual *SATB1* isoforms were plotted. Short isoform uc003cbh.2 and long isoform uc003cbi.2 were used for visualization. Different z-score thresholds were also tested. We reasoned that normalization to the normal tissue samples would bias the results due to possible infiltration of T cells (predominantly expressing *SATB1*).

## 3 Results

### 3.1 Identification of a novel SATB1 protein isoform

SATB1 protein consists of an N-terminal part containing the ULD (ubiquitin-like oligomerization domain; similar in sequence to the PDZ domain) and CUTL (CUT repeat-like) domains which are responsible for its self-oligomerization (Galande et al., 2001; Wang et al., 2012, 2014; Ghosh et al., 2019) and the interaction with other proteins (Yasui et al., 2002; Cai et al., 2003; Notani et al., 2011). The CUT1, CUT2 domains and a homeodomain are responsible for SATB1’s DNA binding activity and are all localized to its C-terminus (**Figure 1a**). Between CUT2 and the homeodomain, there is a glutamine-rich region following the predicted extra peptide (EP) of the long SATB1 isoform [UniProt: long SATB1 isoform E9PVB7_Mouse (795 aa), short SATB1 isoform Q60611 (764 aa)].

In this work, we performed deeply-sequenced stranded-total-RNA-seq experiments in murine thymocytes to identify the existing *Satb1* transcripts. We detected about 5.6-fold enrichment of the transcripts encoding the short (764 amino acids) over the long (795 amino acids) *Satb1* isoform (**Figure 1b**). Note also the presence of seven additional annotated transcripts, three of which may encode for truncated protein isoforms and four untranslated transcripts (do not contain an open reading frame, ORF) which all may have an unknown regulatory function. To validate these findings, we designed primers for RT-qPCR experiments in a way that the forward primer either hybridized at the 3’-end of exon10 and simultaneously at the 5’-end of exon11 (Fwd1) or entirely within the extra exon (Fwd2) of the long isoform, to specifically amplify either the short or the long *Satb1* transcripts, respectively (**Figure 1c**). We confirmed that in murine thymocytes the steady state mRNA levels of the short *Satb1* transcripts were about 3-5 fold more abundant compared to the steady state mRNA levels of the long *Satb1* transcripts (**Figure 1d**). To validate the existence of the long SATB1 isoform, we generated a custom-made antibody specifically targeting the extra peptide of 31 amino acids originating from the extra *Satb1* exon (**Figure 1a**). Moreover, we also developed an antibody targeting the native form of the N-terminal part of SATB1 (the first 330 amino acid residues; **Figure 1a**). The specificity of the custom-made anti-long SATB1 isoform antibody was validated in a heterologous expression system using transient transfection of GFP-tagged SATB1 constructs (**Figure 1e**). It should be noted that none of the commercially available antibodies can distinguish between the two SATB1 isoforms *in vivo*, as they target parts of SATB1 shared by both isoforms. Thus, we can only compare the physiological levels of the long SATB1 isoform to the total SATB1 protein levels. To overcome this limitation and to specifically validate the presence of the long SATB1 protein isoform in primary murine T cells, we designed a serial immunodepletion-based experiment (**Figure 1f**, **Supplementary Figure 1a**). We found that the antibody raised against the extra peptide of the long SATB1 isoform specifically detected only the long SATB1 isoform (**Figure 1g**: upper panel, lane 2; **Supplementary Figure 1b**, lane 2) and not the short SATB1 isoform (**Figure 1g**: upper panel, lane 3; **Supplementary Figure 1b**, upper panel, lane 3). This experiment can also be used for the quantitation of the cellular protein levels of SATB1 isoforms in primary murine thymocytes. Comparison of the two bands for the long (lane 2) and short SATB1 (lane 3) isoform in the lower panel of **Figure 1g** and **Supplementary Figure 1b**, suggested that, upon densitometric quantitation of the band intensity, the long SATB1 isoform protein levels were 1.5× to 2.62× less abundant than the short isoform, according to the two replicates of the immunodepletion experiment, respectively. Moreover, the specificity of the long SATB1 isoform antibody was supported by the fact that it detected no protein in SATB1-depleted T cells (**Supplementary Figure 1c**; *Satb1*^fl/fl^*Cd4*-Cre^+^ knockout mouse – introduced later in the text). Similarly, a validation of the specificity of the anti-all SATB1 isoform custom-made antibody is available (**Supplementary Figure 1d**). The presence of the two isoforms in primary murine thymocytes was also supported by 2D gel electrophoresis coupled to Western blot experiments (**Supplementary Figure 1e**). Additionally, confocal microscopy experiments unraveled the extensive co-localization of the different SATB1 isoforms detected by both types of antibodies (**Supplementary Figure 1f**). Based on this data, we concluded that in murine thymocytes a long SATB1 isoform is expressed at both the mRNA and protein levels.

SATB1 is a nuclear protein and, utilizing confocal microscopy, it was previously shown to display a cage-like pattern in murine thymocytes (Cai et al., 2003). We revisited this question and employed super-resolution microscopy approaches to study SATB1’s subnuclear localization. Our results indicated that the previously described cage-like pattern of SATB1 (Cai et al., 2003) at high resolution, rather resembled individual speckles which may or may not be interconnected (**Figure 1h**). An apparent emerging question, upon detection of these SATB1 speckles, was whether they are involved in the regulation of physiological processes such as transcription.

### 3.2 Co-localization of SATB1 and sites of active transcription

Each super-resolution microscopy approach comes with certain limitations, hence for our applications we compared the results from STED (Stimulated Emission Depletion; provides high resolution, lower risk of bleed-through) with 3D-SIM (3D Structured Illumination Microscopy; allows higher throughput, broader variety of fluorophores and DAPI staining, less photobleaching). In search for functional differences between the two SATB1 isoforms, we probed their subnuclear localization *in vivo* in primary murine thymocytes, utilizing 3D-SIM super-resolution microscopy (**Figure 2a**). The quantification of SATB1 speckles in four nuclear zones, derived based on the relative intensity of DAPI staining, highlighted the localization of SATB1 mainly to the regions with medium to low DAPI staining (zones 3 & 4, **Figures 2a, b**). A similar distribution of the SATB1 signal could also be seen from the fluorocytogram of the pixel-based co-localization analysis between the SATB1 and DAPI signals (**Supplementary Figure 2a**). SATB1’s preference to localize outside heterochromatin regions was supported by its negative correlation with HP1β staining (**Supplementary Figure 2b**). Localization of SATB1 speckles detected by antibodies targeting all SATB1 isoforms or only the long SATB1 isoform, revealed a significant difference in the heterochromatin areas (zone 1, **Figure 2b**), where the long isoform was less frequently present (see also **Figure 2a**). Although, this could indicate a potential difference in localization between the two isoforms, due to the inherent difficulty to distinguish the two, based on antibody staining, we refrain to draw any conclusions.

**Figure 2.**
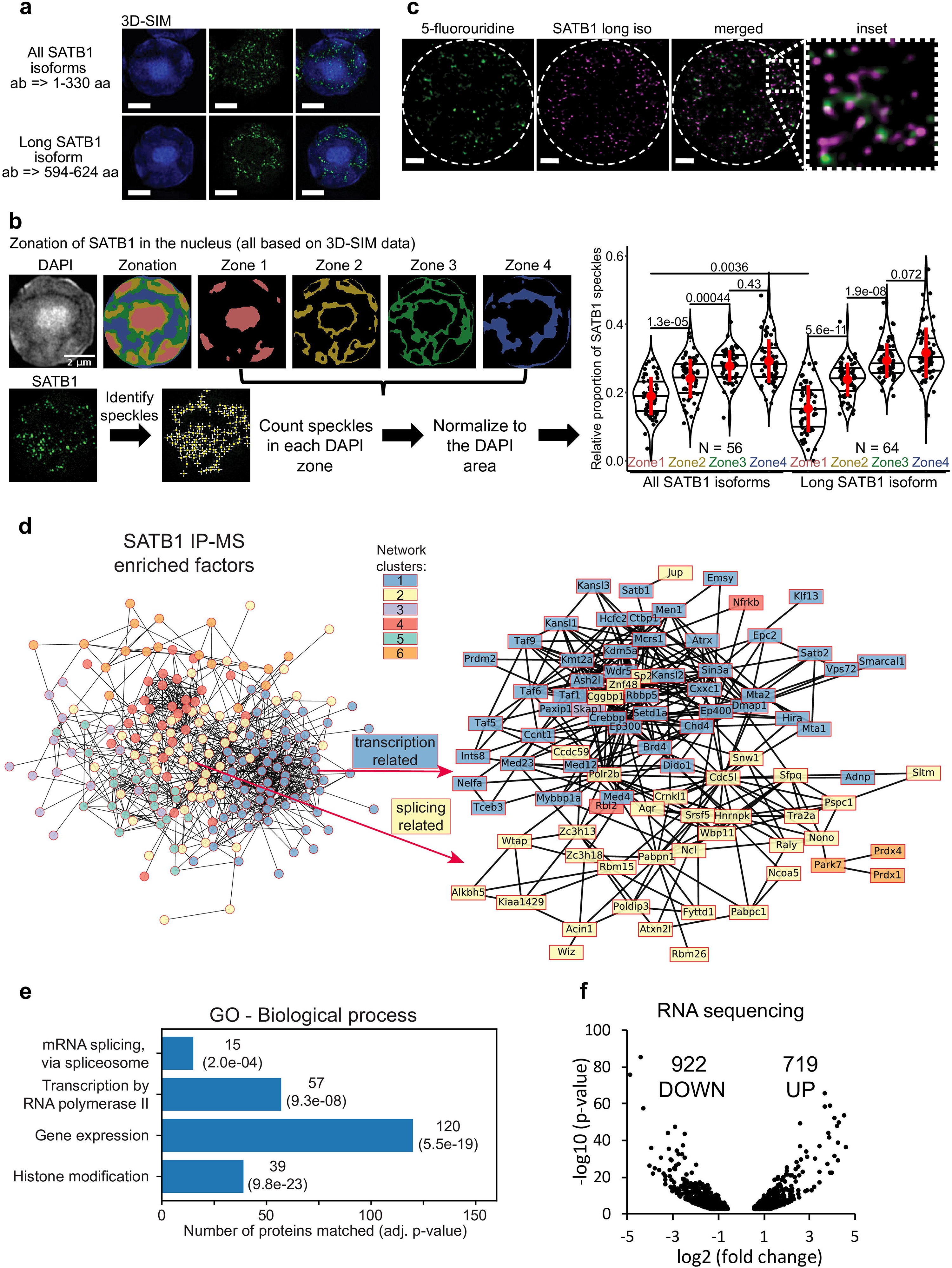
SATB1 is localized in the active nuclear zone and sites of active transcription. **a**, 3D-SIM immunofluorescence experiment utilizing two different SATB1 antibodies to visualize the nuclear localization pattern of the long isoform compared to all SATB1 isoforms. Scale bar 2 µm. **b**, Quantification of the 3D-SIM immunofluorescence images. Nuclei of primary murine thymocytes were categorized into four zones based on the intensity of DAPI staining and SATB1 speckles in each zone were counted. Images used represent a middle z-stack from the 3D-SIM experiments. The graph depicts the differences between long and all SATB1 isoforms’ zonal localization in nuclei of primary murine thymocytes. The horizontal lines inside violin plots represent the 25^th^, 50^th^ and 75^th^ percentiles. Red circles represent the mean ± s.d. *P* values by Wilcoxon rank sum test. **c**, STED microscopy indicating that SATB1 speckles co-localize with sites of active transcription (60-minute pulse 5-fluorouridine treatment). Scale bar 1 µm. See also **Supplementary Figure 3c. d**, STRING network of all significantly enriched SATB1-interacting proteins clustered using the k-means method (left). The full network, including protein names, is available in **Supplementary Figure 4c**. The blue cluster 1 and yellow cluster 2 were enriched for transcription and splicing-related factors, respectively. A more detailed network of selected transcription and splicing-related factors is provided (right). **e**, Gene ontology enrichment analysis of SATB1-interacting proteins revealed factors involved in transcription and splicing. **f**, Volcano plot from stranded-total-RNA-seq experiment displaying the differentially expressed genes (FDR < 0.05) in *Satb1* cKO thymocytes.

The prevailing localization of SATB1 corresponded with the localization of RNA-associated and nuclear scaffold factors, architectural proteins such as CTCF and cohesin, and generally features associated with euchromatin and active transcription (Miron et al., 2020). This was also supported by co-localization of SATB1 with the H3K4me3 histone mark (**Supplementary Figure 2c**), which is known to be associated with transcriptionally active/poised chromatin. Given the localization of SATB1 to the nuclear zones with predicted transcriptional activity (Miron et al., 2020) (**Figure 2b**, zone 3), we investigated the potential association between SATB1 and transcription. We unraveled the localization of SATB1 isoforms and the sites of active transcription as deduced by 5-fluorouridine labeling. Sites of active transcription displayed a significant enrichment in the nuclear zones 3 & 4 (**Supplementary Figure 3a**), similar to SATB1. As detected by fibrillarin staining, SATB1 also co-localized with nucleoli which are associated with active transcription and RNA presence (**Supplementary Figure 3b**). Moreover, we found that the SATB1 signal was in close proximity to nascent transcripts, as detected by STED microscopy (**Figure 2c**). Similarly, the 3D-SIM approach indicated that even SATB1 speckles that appeared not to be in proximity with FU-labeled sites in one z-stack, were found in proximity in another z-stack (**Supplementary Figure 3c**). Additionally, a pixel-based approach to quantify the co-localization of SATB1 with sites of active transcription is presented later in the text in **Figure 3g**, again supporting their co-localization.

**Figure 3.**
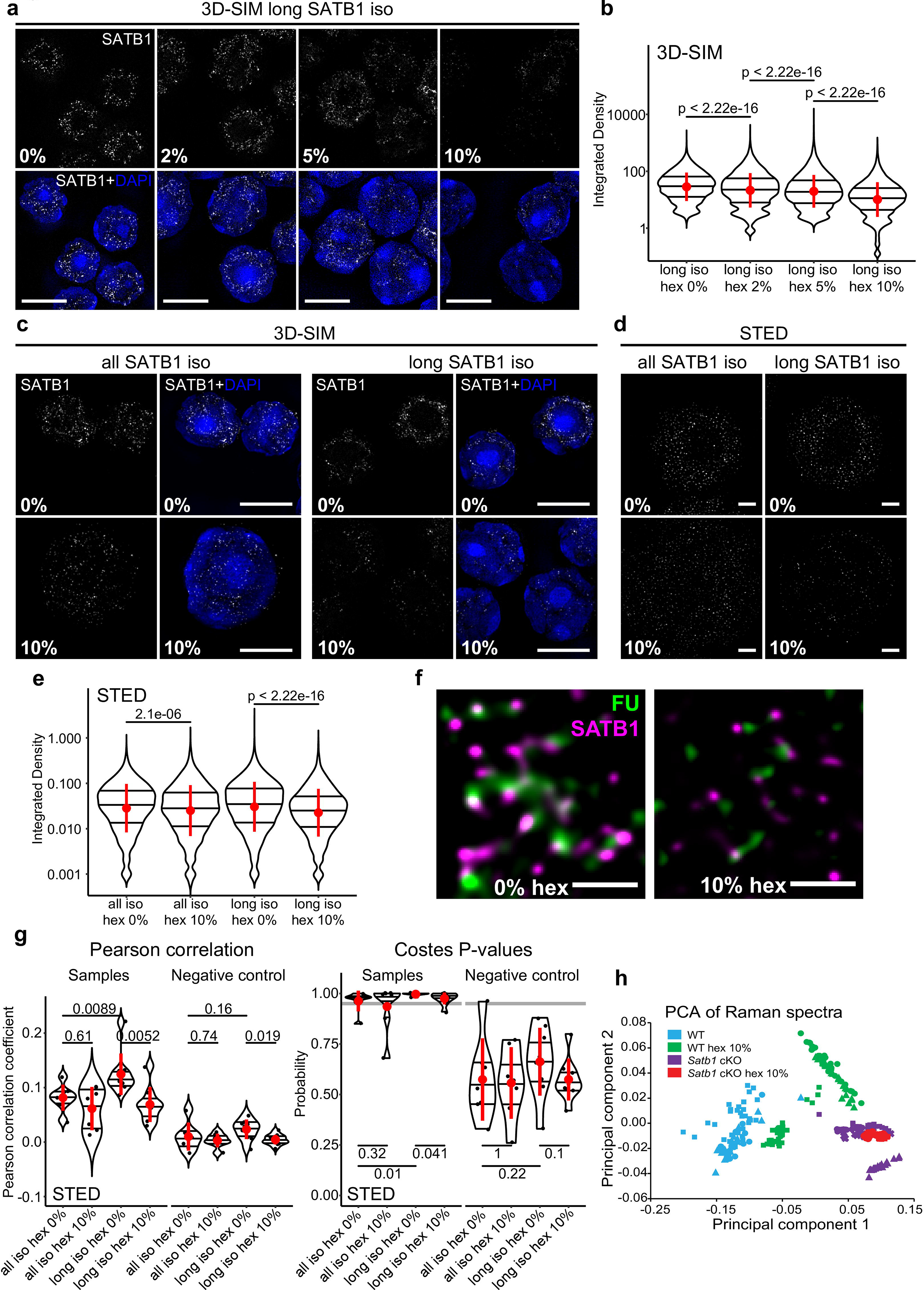
SATB1 forms phase separated droplets *in vivo*. **a**, 3D-SIM immunofluorescence microscopy on 1,6-hexanediol-treated thymocytes. Five minutes treatment with increasing concentrations of 1,6-hexanediol gradually solubilized long SATB1 isoform speckles. **b**, Quantification of results in **a** showing a gradual decrease of SATB1 signal, indicating its sensitivity to 1,6-hexanediol treatment. **c**, Comparison of immunofluorescence signal based on the antibody targeting all the SATB1 isoforms (Santa Cruz Biotechnology, sc-5990) and only the long isoform (Davids Biotechnology, custom-made) upon 1,6-hexanediol treatment, detected by 3D-SIM microscopy. **d**, Immunofluorescence experiment as indicated in **c** but detected by STED microscopy. Scale bar 0.5 µm. **e**, Quantification of results in **d** showing a more dramatic decrease of long SATB1 isoform signal upon 1,6-hexanediol treatment compared to the all SATB1 isoforms staining. **f**, Co-localization of SATB1 and fluorouridine-stained sites of active transcription (inset visualized in Figure 2c) and its deregulation upon 10% 1,6-hexanediol treatment detected by STED microscopy. Scale bar 0.5 µm. **g**, LEFT: Pearson correlation coefficients (PCC) derived from pixel-based co-localization analysis between long and/or all SATB1 isoforms and fluorouridine-stained sites of active transcription, with and without 10% 1,6-hexanediol treatment. RIGHT: the Costes P-values (Costes et al., 2004) derived from randomly shuffled chunks of analyzed images (100 randomizations). The grey line indicates a 0.95 level of significance. Images used for the analysis were generated by STED microscopy. Complete results of the co-localization analysis for both STED and 3D-SIM experiments, including Manders’ coefficients are depicted in **Supplementary Figures 5a-c**. **h**, Principal component analysis of Raman spectra from WT and *Satb1* cKO thymocytes with and without 10% 1,6-hexanediol treatment. Each point represents measurements from an individual cell. For each condition, 2-5 biological replicates were used. See the extracted Raman spectra of the two main principal components that were used to cluster the data in **Supplementary Figure 5d**. In **b**, **e** and **g**, the horizontal lines inside violin plots represent the 25^th^, 50^th^ and 75^th^ percentile. Red circles represent the mean ± s.d. *P* values by Wilcoxon rank sum test.

We further performed co-immunoprecipitation experiments (utilizing an antibody targeting all SATB1 protein isoforms) coupled to mass spectrometry analysis (IP-MS), to determine SATB1’s interactome in murine thymocytes (**Supplementary Figure 4a-b**; **Supplementary File 1**). Apart from subunits of chromatin modifying complexes that were also detected in previous reports (Yasui et al., 2002; Fujii et al., 2003; Kumar et al., 2006; Purbey et al., 2009; Notani et al., 2010), unbiased k-means clustering of the significantly enriched SATB1 interactors revealed two major clusters consisting mostly of proteins involved in transcription (blue cluster 1; **Figure 2d** and **Supplementary Figure 4c**) and splicing (yellow cluster 2; **Figure 2d** and **Supplementary Figure 4c**). Both transcription and splicing were also significantly enriched terms in the gene ontology enrichment analysis of SATB1 interactors (**Figure 2e**; **Supplementary File 2**). These findings thus supported our hypothesis regarding SATB1’s involvement in thymocyte RNA production and processing as inferred from its nuclear localization.

To further investigate the functional link between SATB1 and gene transcription and splicing, we performed stranded-total-RNA-seq experiments in wild type (WT) and *Satb1*^fl/fl^*Cd4*-Cre^+^ (*Satb1* cKO) murine thymocytes, with SATB1 being specifically depleted at the CD4^+^CD8^+^ T cells during thymocyte development (Zelenka et al., 2022). *Satb1* cKO animals display severely impaired T cell development associated with largely deregulated transcriptional programs as previously documented (Kakugawa et al., 2017; Kitagawa et al., 2017; Zelenka et al., 2022). We have previously shown (Zelenka et al., 2022) that long SATB1 isoform-specific binding sites (GSE173446; Zelenka et al., 2022) were associated with increased chromatin accessibility, with a visible drop in chromatin accessibility in *Satb1* cKO. Moreover, the drop in chromatin accessibility was especially evident at the transcription start site of genes, suggesting that the long SATB1 isoform is directly involved in transcriptional regulation. Consistent with these findings and with SATB1’s nuclear localization at sites of active transcription, we identified a vast transcriptional deregulation in *Satb1* cKO with 1,641 (922 down-regulated, 719 up-regulated) differentially expressed genes (**Figure 2f**). Additionally, there were 2,014 genes with altered splicing efficiency (**Supplementary Figure 4d-e**; **Supplementary Files 3-4**). We should also note that the extent of splicing deregulation was directly correlated with long SATB1 isoform binding (**Supplementary Figure 4d**).

In summary, we found that in primary thymocyte nuclei, SATB1 localized in euchromatin and interchromatin regions in close proximity to actively transcribed genes, interacted with protein complexes involved in the regulation of gene transcription and mRNA splicing and upon *Satb1* deletion, both physiological processes of transcription and splicing were deregulated.

### 3.3 Liquid-like states of SATB1 is primary T cells

SATB1 has a predicted phase separation potential (Wang et al., 2018). Here we demonstrated its connection to transcription and found that it forms spherical speckles (**Figure 1h**), markedly resembling phase separated transcriptional condensates (Cho et al., 2018; Sabari et al., 2018). Thus, we next investigated whether the observed nuclear SATB1 speckles could be a result of LLPS. We should note that since we were primarily focused on studying the physiological roles of SATB1 in primary murine T cells, we were limited by the experimental options available. To probe differences in phase separation in murine primary cells, without any intervention to SATB1 structure and expression, we first utilized 1,6-hexanediol treatment, which was previously shown to dissolve liquid-like droplets (Kroschwald et al., 2017). If SATB1 was indeed undergoing phase separation into the transcriptional condensates, we would expect to detect a sensitivity of the SATB1 speckles to the hexanediol treatment, followed by decreased co-localization with fluorouridine-labeled sites of active transcription. The long SATB1 isoform speckles displayed such sensitivity as demonstrated by a titration series with increasing concentrations of 1,6-hexanediol treatment followed by 3D-SIM super-resolution microscopy analysis (**Figure 3a-b**; note the log scale). Moreover, analysis with 3D-SIM microscopy revealed that the long SATB1 isoform was more sensitive to hexanediol treatment than collectively all SATB1 protein isoforms (**Figure 3c**). This was also supported by data from STED microscopy (**Figure 3d**) as quantified in **Figure 3e**. Additionally, hexanediol treatment highly decreased co-localization between SATB1 and fluorouridine-labeled sites of active transcription (**Figure 3f**). To quantify the difference in co-localization between SATB1 and fluorouridine-labeled sites of active transcription, as well as the changes upon hexanediol treatment, we used a pixel-based co-localization analysis (**Figure 3g** and **Supplementary Figure 5a-c**). Based on these results, the long SATB1 isoform displayed a higher association with sites of active transcription and this association was more sensitive to the hexanediol treatment than collectively all SATB1 isoforms.

To expand our study regarding the effect of 1,6-hexanediol in dissociating SATB1 speckles in primary T cells, we employed Raman spectroscopy, a non-invasive label-free approach, which is able to detect changes in chemical bonding. Raman spectroscopy was already used in many biological studies, such as to predict global transcriptomic profiles from living cells (Kobayashi-Kirschvink et al., 2018) and also in research of protein LLPS and aggregation (Murthy et al., 2019; Banc et al., 2021; Agarwal et al., 2022; Shuster and Lee, 2022; Yokosawa et al., 2022). Thus, we reasoned that it may also be used to study phase separation in primary T cells. We measured Raman spectra in primary thymocytes derived from both WT and *Satb1* cKO animals and compared them with spectra from cells upon 1,6-hexanediol treatment. Principal component analysis of the resulting Raman spectra (**Supplementary Figure 5d**), clustered the treated and non-treated *Satb1* cKO cells together, while the WT cells clustered separately (**Figure 3h**). The WT 1,6-hexanediol treated cells clustered between the aforementioned groups. These results suggested that the principal component 1 (PC1) could discriminate WT and *Satb1* cKO cells based on chemical bonding provided by SATB1 protein. These bonds were probably enriched for weak interactions responsible for phase separation that are susceptible to hexanediol treatment. This shifted the cluster of WT treated cells towards the *Satb1* cKO cells. However, the remaining covalent bonds differentiated the WT samples from *Satb1* cKO cells where all bonds provided by SATB1 were absent. This approach demonstrated that Raman spectroscopy can be utilized as a non-invasive research approach to unravel the global chemical state of a cell and its changes upon various conditions. Moreover, together with the light microscopy experiments, it allowed us to investigate SATB1’s biophysical behavior *in vivo*.

To further test our hypothesis that SATB1 can undergo phase separation, we have conducted turbidity assays, showing positive correlation between absorbance and increasing concentration of recombinant SATB1 (**Supplementary Figure 6a**). Similarly, SATB1 displayed increased absorbance with time (**Supplementary Figure 6b**), which was further intensified upon addition of its DNA target region RHS6 (**Supplementary Figure 6c**). These data were additionally supported by centrifugation assays (**Supplementary Figure 6d**), collectively suggesting that SATB1 can undergo phase separation.

### 3.4 The long SATB1 isoform is more prone to aggregation *in vitro*

To further investigate the phase separation capacity of SATB1, we utilized the light-inducible CRY2-mCherry optoDroplet system (Shin et al., 2017) (**Figure 4a**). This tool allows studying the phase separating properties of individual parts of a protein in a controlled manner. The CRY2-mCherry chimeric constructs, encoding part of a protein of interest are transiently transfected into cells. Following illumination by high energy light, the photo-responsive CRY2 protein is induced to homo-oligomerize while bringing the protein molecules of interest into spatial proximity. Only proteins with phase separating properties will then form droplets, which are visualized by accumulation of the mCherry signal. First, we selected the multivalent N-terminal part of SATB1 (1-298 aa), which is known for its involvement in protein-protein interactions (Galande et al., 2001; Yasui et al., 2002; Cai et al., 2003; Notani et al., 2011; Wang et al., 2012, 2014). Note, that the multivalent nature of a protein indicates its predisposition to undergo LLPS (Banani et al., 2017; Shin and Brangwynne, 2017; Alberti et al., 2019). Moreover, the selected region of SATB1 includes part of a prion-like domain (**Figure 4a**), which makes the N-terminus of SATB1 a prospective candidate with phase separation potential. Transiently transfected NIH-3T3 cells were activated with a 488 nm laser, every two seconds, to initiate CRY2 association and mCherry signal was detected. For the analysis, cells were grouped into two categories based on the original mCherry relative fluorescence, reflecting the transfection efficiency, as the protein concentration directly affects the speed of droplet assembly (Shin et al., 2017). The construct encompassing the N-terminus of SATB1 formed optoDroplets at a similar or even faster rate than the FUS positive control (**Figure 4b**, live videos in **Supplementary File 5**), a popular model for LLPS studies (Shin et al., 2017). The low SATB1 concentration group of cells formed the first optoDroplets ∼10 seconds upon induction, compared to FUS where the first optoDroplets appeared after ∼14-20 seconds upon stimulation. Cells with higher SATB1 expression levels displayed even faster optoDroplet assembly, where both FUS and the N-terminus of SATB1 started forming optoDroplets at ∼6 seconds upon activation. The formed speckles resembled phase separated droplets based on their round shape and ability to fuse (**Figure 4c** and **Supplementary Figure 7a**). The droplet assembly rate for both the N-terminal part of SATB1 and FUS proteins (**Figure 4d**) highlighted the direct correlation between protein concentration and the speed of droplet assembly. We detected a higher number of FUS optoDroplets that were formed, however the N-terminus SATB1 optoDroplets grew in bigger sizes (**Figure 4b** and **Supplementary Figure 7b**). FUS optoDroplets disassembled entirely after 100 and 250 seconds of inactivation time (488 nm laser off), for the low and high protein expression groups, respectively. In contrary, the optoDroplets formed by the N-terminus of SATB1 displayed a much slower disassembly rate and often even after ten minutes of inactivation time there were some rigid structures still present (**Figure 4e**). The group of cells with a lower concentration of the SATB1 N-terminus displayed a gradual decrease in droplet size during inactivation time, unlike the high concentration group (**Supplementary Figure 7c**). Following the original faster assembly of SATB1 structures, they also turned more quickly into long-lasting aggregates. Moreover, the high concentration group displayed decreased circularity of droplets compared to the low concentration samples (**Supplementary Figure 7a**). FRAP (fluorescence recovery after photobleaching) experiments indicated that SATB1 droplets had around 43% of an immobile fraction compared to only 17% for FUS (**Figure 4f**). However, the mobile fraction of SATB1 droplets manifested a halftime of recovery 3.3 seconds, much lower than the halftime of 11.4 seconds of FUS droplets. These results suggested that the N-terminus constructs could have been generally over-activated. However, taken all our data together, we reasoned that SATB1 does not behave like common phase separating proteins. The N-terminus of SATB1 is shared by both the short and long SATB1 isoforms, so we next aimed to compare the behavior of full length SATB1 isoforms. In transient transfection experiments of EGFP-SATB1 constructs, without the photo-responsive CRY2 protein, we could detect similar FRAP kinetics for the short and long full length SATB1 isoforms (**Supplementary Figure 7f**). Next, we again utilized the optoDroplet system – to investigate whether the two full length SATB1 isoforms could undergo phase separation similar to the shorter N-terminal part. Upon activation, both full-length isoforms displayed visible ultrastructural changes which were reversible upon inactivation (**Figure 5a-b**). However, these did not resemble the optoDroplets detected for FUS and the N-terminus of SATB1, as deduced from the number and size of the optoDroplets detected (**Supplementary Figure 7d-e**). Although the full length SATB1 bears a nuclear localization signal (aa 20-40), we identified a fraction of cells where the protein was localized in the cytoplasm (**Figure 5c**). The mislocalized long SATB1 isoform protein was often found in a form of rigid structures resembling protein aggregates, unlike the short SATB1 isoform (**Figure 5c**). Computational analysis, using the algorithm *cat*GRANULE (Bolognesi et al., 2016), of the protein sequence for both murine SATB1 isoforms indicated a higher propensity of the long SATB1 isoform to undergo LLPS with a propensity score of 0.390, compared to 0.379 for the short isoform (**Figure 5d**). This difference was dependent on the extra peptide of the long isoform. Out of the 31 amino acids comprising the murine extra peptide, there are six prolines, five serines and three glycines – all of which contribute to the low complexity of the peptide region (Shin and Brangwynne, 2017) (**Figure 5e**). Moreover, the extra peptide of the long SATB1 isoform directly extends the PrLD and IDR regions as indicated in **Figure 4a**. Since protein aggregation has been previously described for proteins containing poly-Q domains and PrLDs (Williams and Paulson, 2008; March et al., 2016; Harrison and Shorter, 2017; Peskett et al., 2018), we next generated truncated SATB1 constructs encoding two of its IDR regions, the PrLD and poly-Q domain and in the case of the long SATB1 isoform also the extra peptide neighboring the poly-Q domain (**Figure 1a** and **Figure 4a**). The constructs containing the extra peptide of the long SATB1 isoform frequently resulted in protein aggregates which did not respond to CRY2 activation (**Figure 5f**). The constructs derived from the short SATB1 isoform did not form almost any aggregates or only in a very small fraction of cells (**Figure 5g**). Our data indicate the potential of SATB1 to undergo phase transitions leading to the formation of aggregated structures. The two SATB1 isoforms differ in their propensity to undergo these transitions and to aggregate, which may constitute a crucial determinant of their differential biological roles *in vivo*.

**Figure 4.**
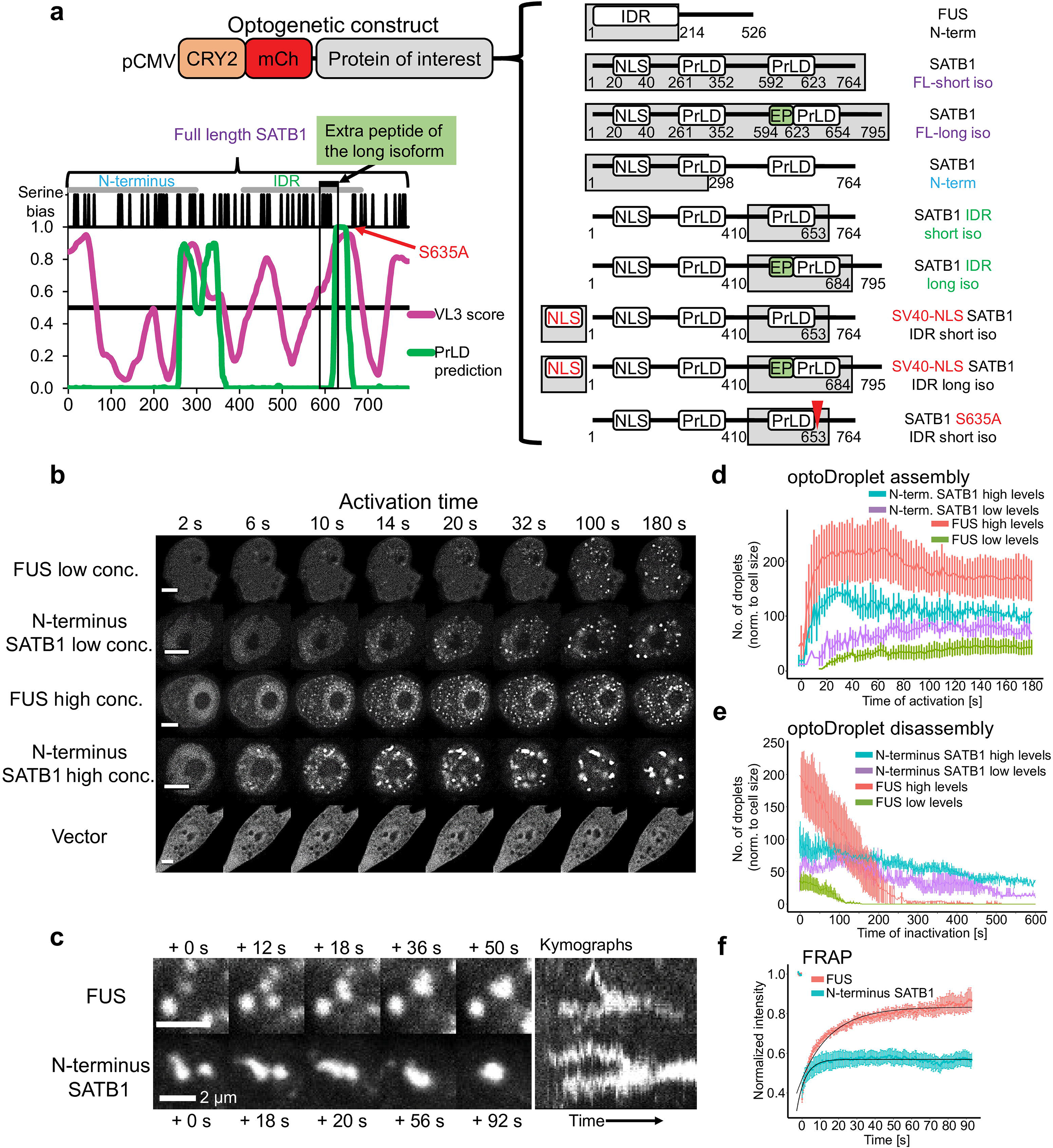
The N-terminus of SATB1 forms phase separated droplets. **a**, Graphical representation of all CRY2-mCherry constructs generated. The graph displays the IDR prediction by VL3 PONDR score in magenta and the prediction of prion-like domains by PLAAC algorithm in green. **b**, Formation of optoDroplets upon 488 nm pulse illumination in live cells transiently transfected with CRY2-mCherry-N-terminus_SATB1 constructs as well as with negative (vector) and positive control constructs (N-terminal part of FUS protein). Two groups of cells were identified based on the original relative expression levels of the recombinant proteins. Scale bar 5 µm. **c**, OptoDroplets displayed a round shape and the ability to coalesce. **d**, Number of optoDroplets normalized to the cell size at each time point upon activation, to visualize the speed of droplet assembly and its dependence on the original concentration of the protein. **e**, Number of optoDroplets normalized to the cell size at each time point of inactivation (488 nm laser was off) to visualize the speed of droplet disassembly and its dependence on the original concentration of the protein. **f**, FRAP experiment utilizing either the N-terminus of SATB1 construct or the FUS positive control construct. The fitted curves were generated by the exponential recovery curve fitting function of Fiji. In **d**, **e** and **f**, the error bars represent the s.e.m.

**Figure 5.**
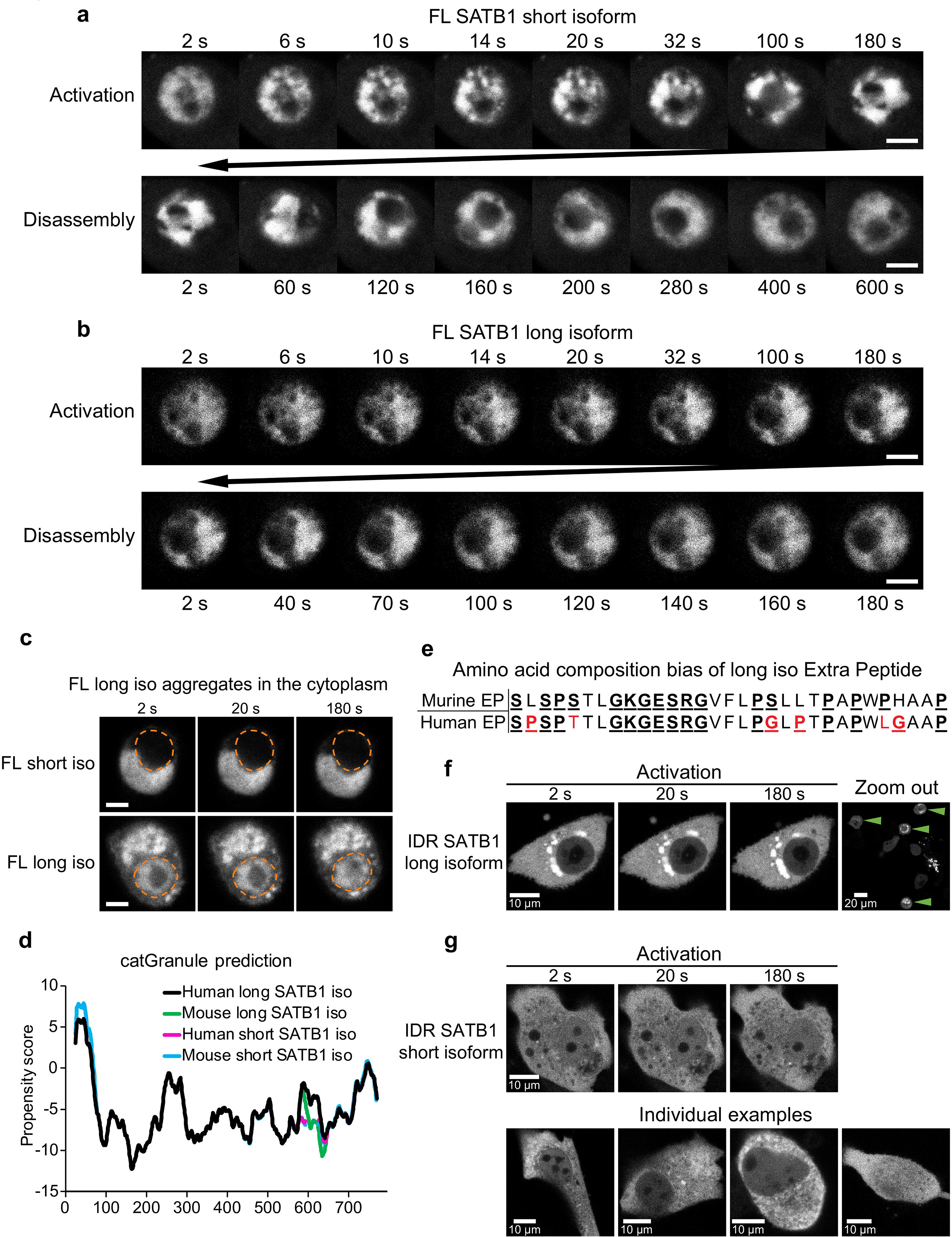
Discrete biophysical properties of SATB1 protein isoforms. **a**, CRY2-mCherry constructs expressing full length short SATB1 isoform underwent reversible ultrastructural changes. Scale bar 5 µm. **b**, The same experiment as in **a** but capturing CRY2-mCherry constructs expressing full length long SATB1 isoform. Scale bar 5 µm. **c**, In a fraction of cells, the full-length SATB1 constructs were mislocalized outside the nucleus. For constructs with the long SATB1 isoform this triggered formation of irreversible aggregates. Scale bar 5 µm. **d**, Comparison of LLPS propensity predicted by the *cat*GRANULE (Bolognesi et al., 2016) algorithm of human (Q01826-1 and Q01826-2, for short and long isoform, respectively; UniProtKB) and murine (Q60611 and E9PVB7, for short and long isoform, respectively; UniProtKB) SATB1 protein isoforms. Human long SATB1 isoform displays higher phase separation propensity compared to murine protein. **e**, Both murine and human long SATB1 isoforms have an increased propensity to undergo phase separation compared to short isoforms due to the presence of the extra peptide with a compositional bias (enrichment in S, G, Q, P, E, K and R amino acids in the murine SATB1). Red letters indicate amino acid differences between murine and human peptide. **f**, CRY2-mCherry constructs encompassing only the two SATB1 IDR regions and the poly-Q domain (as indicated in Figure 4a and Figure 1a), but not the NLS signal, displayed much higher rates of cytoplasmic aggregation for the long SATB1 isoform compared to the short isoform. Green arrows indicate cells with long SATB1 isoform aggregates. **g**, The same experiment as in **f**, but for the short SATB1 isoform IDR constructs.

### 3.5 Modes of regulation of SATB1 phase transitions

To determine why SATB1 forms aggregates in the cytoplasm, we next investigated the potential means of regulation for the SATB1 phase transitions. Phase separating properties of prion-like proteins are regulated by the presence of nuclear RNA, which keeps them in a soluble form when in the nucleus (Maharana et al., 2018). However, currently it is not known whether SATB1 interacts with RNA at all. The data presented here involve SATB1 to both transcription and splicing (**Figure 2** and **Figure 3**). Moreover, SATB1 was originally discovered as a protein of the nuclear matrix (Dickinson et al., 1992), where RNA represents one of its main components and thus implying that SATB1’s association with RNA is plausible. To test this hypothesis, we performed nuclear matrix extraction experiments, including RNase A treatment, to investigate the potential association of SATB1 with RNA. A fraction of SATB1 remained in the nucleus even after the extraction of cells with high concentration of salt or treatment with DNase I, though the treatment with RNase A alone depleted SATB1 from the cell nucleus of murine thymocytes (**Figure 6a**). One of the prominent RNAs associated with the nuclear matrix is lncRNA Xist, whose interaction with SATB1 was already predicted (Agostini et al., 2013). To further validate these findings, we performed RNA immunoprecipitation experiments utilizing antibodies against the SATB1 protein and we detected the association between SATB1 and Xist lncRNA (**Figure 6b**). Moreover, when we forced localization of the long SATB1 isoform IDR constructs to the nucleus, by fusing them to the SV40 nuclear localization signal, the protein signal remained diffused in the cell nucleus (**Figure 6c**) and notably it did not respond to the CRY2 activation even after 20 minutes of activation. Therefore, here we demonstrated that SATB1 in primary murine T cells is capable of association with RNA, that its localization in the nucleus is highly dependent on the presence of nuclear RNA and that the nuclear environment affects phase transitions of SATB1.

**Figure 6.**
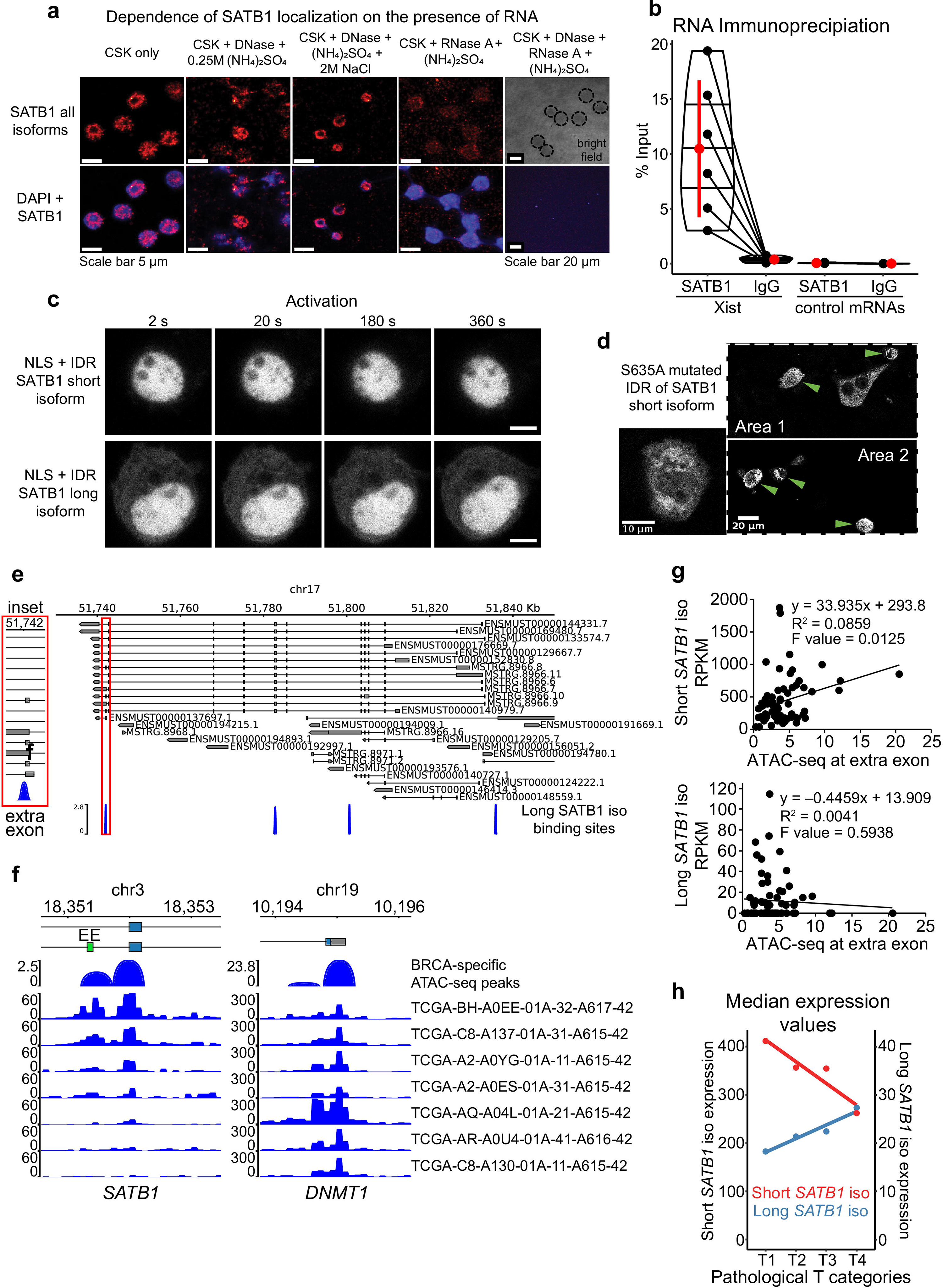
Modes of regulation of SATB1 phase transitions. **a**, Nuclear matrix extraction coupled to immunofluorescence analysis revealed that a fraction of SATB1 protein (Santa Cruz Biotechnology, sc-5990) remained in the cell nucleus even after DNase I treatment and high-salt extraction. However, RNase A treatment almost completely depleted SATB1 from the nucleus, indicating a high level of association between SATB1 and nuclear RNA. **b**, SATB1 co-immunoprecipitation confirmed the association of SATB1 with lncRNA Xist. Black dots represent individual % input measurements for Xist RIP (3 biological replicates, 4 Xist regions targeted) and control RIP experiments (3 biological replicates; 3 control RNAs: *Tbp*, *Smc1a*, Dxz4). The horizontal lines inside violin plots represent the 25^th^, 50^th^ and 75^th^ percentiles. Red circle represents the mean ± s.d. Mean fold enrichment for Xist RIP is 36 ± 20 s.d. **c**, Forced nuclear localization of CRY2-mCherry constructs harboring the short or the long SATB1 isoform IDR and the SV40 NLS displayed no protein association in response to 488 nm laser activation. Scale bar 5 µm. **d**, The S635A mutation, preventing phosphorylation of the short SATB1 isoform highly increased the aggregation propensity of the corresponding CRY2-mCherry-IDR construct. Green arrows indicate cells with the SATB1 aggregates. **e**, Murine *Satb1* transcripts reconstructed based on deeply-sequenced stranded-total-RNA-seq data. Both long and short *Satb1* isoforms were produced from multiple promoters. The long SATB1 isoform binding sites were retrieved from GSE173446 (Zelenka et al., 2022). **f**, Human TCGA breast cancer (BRCA) patient-specific ATAC-seq peaks (Corces et al., 2018) span the extra exon (EE; labeled in green) of the long *SATB1* isoform. Note the differential chromatin accessibility in seven selected patients, emphasizing the heterogeneity of *SATB1* chromatin accessibility in cancer. Chromatin accessibility at the promoter of housekeeping gene *DNMT1* is shown as a control. **g**, Increased ATAC-seq signal in human breast cancer patients is positively correlated with the expression of the short *SATB1* isoform and negatively correlated with the long isoform expression. **h**, In human breast cancer patients, high pathological T categories (indicating bigger extent of the primary tumor, presumably indicating worse prognosis; labeling based on the TNM cancer staging system) were associated with higher expression of the long *SATB1* isoform. In contrast, the expression of the short *SATB1* isoform was negatively correlated with the pathological T categories. Median RNA expression values for both isoforms based on all transcripts with non-zero expression values are displayed. Red circles represent the median values.

Another factor affecting the phase transitions of proteins includes their post-translational modifications. More specifically, the phosphorylation of the PrLD of FUS protein was shown to prevent its aggregation (Monahan et al., 2017; Ding et al., 2020). Similarly, the efficient disassembly of stress granules and other membraneless organelles was dependent on phosphorylation (Wippich et al., 2013, 3; Rai et al., 2018; Shattuck et al., 2019). SATB1 harbors many verified and predicted phosphorylation sites (Zelenka and Spilianakis, 2020). The selected IDR region of SATB1 (**Figure 4a**) contains multiple serine residues which are targets of phosphorylation. Based on DNA affinity chromatography coupled to mass spectrometry, we identified a novel SATB1 phosphorylation site, S635 (S666 in the long isoform, **Supplementary Figure 8a**). First, we performed DNA affinity purification assay to demonstrate the importance of phosphorylation for SATB1 binding to DNA (**Supplementary Figure 8b**). Upon phosphatase treatment, SATB1 could not be captured and pulled-down by a biotinylated RHS6 DNA probe (RAD50 DNase I hypersensitive site of the TH2 Locus Control Region (Spilianakis et al., 2005; Williams et al., 2013)), where it effectively binds when phosphatase inhibitors are added (**Supplementary Figure 8c**). To further investigate the impact of phosphorylation in position S635, which directly follows the PrLD, we generated a S635A SATB1 mutant, abolishing phosphorylation at this site (**Figure 4a**). We performed DNA affinity purification assays, utilizing protein extracts from transiently transfected cells (no endogenous SATB1 expressed) with either the wild type or mutated SATB1 and found that this mutant had abrogated DNA binding ability (**Supplementary Figure 8d**). Even more importantly, a CRY2-mCherry construct with the mutated part (S635A) of the short SATB1 isoform developed aggregates in the cytoplasm (**Figure 6d**), similar to those of the cytoplasmic long isoform (**Figure 5c**). This observation suggested that phosphorylation of SATB1 plays an important role in its phase transitions, affecting DNA binding and the potential regulatory effects of SATB1 on genome organization and transcriptional gene regulation. In relation to this, a functional association between SATB1 and PML bodies was already described in Jurkat cells (Kumar et al., 2007). We should note that PML bodies represent an example of phase separated nuclear bodies (Banani et al., 2016) associated with SATB1. Targeting of SATB1 into PML bodies depends on its phosphorylation status; when phosphorylated it remains diffused, whereas unphosphorylated SATB1 is localized to PML bodies (Tan et al., 2010). This is in line with the phase separation model as well as with our results from S635A mutated SATB1, which has a phosphorylation blockade promoting its phase transitions and inducing aggregation. To further test whether SATB1 dynamics is affected by its association with PML, we co-transfected short and long full length SATB1 isoforms with PML isoform IV. The dynamics of long SATB1 isoform was affected more dramatically by the association with PML compared to the short SATB1 isoform (**Supplementary Figure 8e**), which again supports the differential behavior of the two SATB1 isoforms.

Furthermore, a regulatory mechanism coordinating the production of the short and long *Satb1* isoforms could also control SATB1’s phase transitions – given their differential calculated LLPS propensity. Such a mechanism could rely on the differential promoter usage as recently reported (Khare et al., 2019; Patta et al., 2020). Our stranded-total-RNA-seq experiments revealed a series of long isoform transcripts originating from various *Satb1* promoters, though indistinguishable from promoters priming the short isoforms (**Figure 6e**). Therefore, we reasoned that a more plausible hypothesis would be based on the regulation of alternative splicing.

We have already reported that the long SATB1 isoform binding sites display increased chromatin accessibility and this is dropped in the *Satb1* cKO, collectively indicating that long SATB1 isoform binding promotes increased chromatin accessibility (Zelenka et al., 2022). We identified a binding site specific to the long SATB1 isoform right at the extra exon of the long isoform (**Figure 6e**). Moreover, the study of alternative splicing based on our RNA-seq analysis revealed a deregulation in the usage of the extra exon of the long *Satb1* isoform (the only *Satb1* exon affected) in *Satb1* cKO cells (deltaPsi = 0.12, probability = 0.974; **Supplementary File 4**). These data suggest that SATB1 itself is able to control the levels of the short and long *Satb1* isoforms. Following the connection between SATB1 binding and chromatin accessibility, a possible mechanism controlling the alternative splicing of *Satb1* gene is based on its kinetic coupling with transcription. Several studies indicated how histone acetylation and generally increased chromatin accessibility may lead to exon skipping, due to enhanced RNA polymerase II elongation (Schor et al., 2009; Zhou et al., 2011). Thus, the increased chromatin accessibility promoted by the long SATB1 isoform binding at the extra exon of the long isoform, would increase RNA polymerase II read-through leading to decreased time available to splice-in the extra exon and thus favoring the production of the short SATB1 isoform in a negative feedback loop manner. This potential regulatory mechanism of SATB1 isoform production is supported by the increased usage of the extra exon in the absence of SATB1 in *Satb1* cKO (**Supplementary File 4**). To further address this, we utilized the TCGA breast cancer dataset (BRCA) as a system expressing SATB1 (Han et al., 2008). ATAC-seq experiments for a series of human patients with aggressive breast cancer (Corces et al., 2018) revealed differences in chromatin accessibility at the extra exon of the *SATB1* gene (**Figure 6f**). In line with the “kinetic coupling” model of alternative splicing, the increased chromatin accessibility at the extra exon (allowing faster read-through by RNA polymerase) was positively correlated with the expression of the short *SATB1* isoform and negatively correlated with expression of the long *SATB1* isoform (**Figure 6g**). Moreover, we investigated whether the differential expression of *SATB1* isoforms was associated with poor disease prognosis. Worse pathological stages of breast cancer and expression of *SATB1* isoforms displayed a positive correlation for the long isoform but not for the short isoform (**Figure 6h** and **Supplementary Figure 9a**). This was further supported by worse survival of patients with increased levels of long *SATB1* isoform and low levels of estrogen receptor (**Supplementary Figure 9b**). Overall, these observations not only supported the existence of the long *SATB1* isoform in humans, but they also shed light at the potential link between the regulation of *SATB1* isoforms production and their involvement in pathological conditions.

## 4 Discussion

In this work we characterized a novel isoform of SATB1 protein, an important transcription factor and genome organizer implicated in intra-thymic T cell development. We demonstrated that the long SATB1 isoform is expressed at high levels in all SATB1-expressing cells. SATB1 in primary murine thymocytes was mainly localized in the transcriptionally active nuclear zone and co-localized with sites of active transcription. Correspondingly, SATB1 depletion in CD4 T cells resulted in deregulated transcriptional programs as also previously demonstrated (Kakugawa et al., 2017; Kitagawa et al., 2017; Zelenka et al., 2022) and in altered splicing efficiency. Moreover, the long SATB1 isoform-specific speckles displayed higher co-localization with fluorouridine-labeled sites of active transcription, compared to the speckles detected by antibodies targeting collectively all SATB1 isoforms, suggesting that the long isoform may be more frequently involved in the regulation of transcription compared to the short isoform.

Moreover, the long isoform speckles in primary T cells displayed higher susceptibility to hexanediol treatment, suggesting potential differences in the phase separation propensity of the two isoforms. To address these questions, we used a heterologous system with genetically engineered optogenetic constructs probing different SATB1 polypeptides. The N-terminus of SATB1, shared by both SATB1 isoforms, displayed rapid formation of liquid-like droplets in the CRY2-mCherry optoDroplet system. We analyzed transiently transfected cells with varying concentration of the recombinant protein. Notably the low-expressing cells transfected with the N-terminus of SATB1 displayed even faster droplet assembly compared to the positive control FUS. Correspondingly, in the highly-expressing cells, SATB1 tended to form larger structures, generally displaying signs of over-saturation and thus conversion into gel-like or solid-like structures (Alberti et al., 2019), known for PrLD proteins. SATB1 contains two PrLD regions, one of which is included at the 3’ end of the SATB1 N-terminal construct. This PrLD also likely facilitated formation of liquid-like optoDroplets of the N-terminus SATB1 constructs. The second PrLD directly extends the region harboring the extra peptide of the long SATB1 isoform, which by itself contributed to the higher theoretical LLPS propensity of the long isoform and its aggregation in the cytoplasm. However, the forced nuclear localization of the second PrLD with the extra peptide of the long isoform, even at very high concentrations, did not indicate any optoDroplet formation. This suggested that the N-terminal part of SATB1 with its ability to form multivalent interactions was necessary for SATB1’s phase transitions. Although our data indicated the differential biophysical behavior of the two SATB1 isoforms, due to limited experimental approaches and main focus on primary T cells, we refrain to explicitly state whether SATB1 speckles are formed by LLPS, polymer-polymer phase separation or by other mechanisms (Erdel and Rippe, 2018).

We next showed that the nuclear RNA environment contributes to the solubility of nuclear SATB1, resulting in protein aggregation when mislocalized into the cytoplasm. Apart from nuclear RNA, buffering a protein’s phase transitions (Maharana et al., 2018), two studies described the role of nuclear-import receptors in chaperoning and disaggregating aberrant phases of PrLD containing proteins (Guo et al., 2018; Yoshizawa et al., 2018). The latter does not seem to apply to SATB1, as SATB1 possesses a unique NLS lacking the clusters of basic residues, thus being dissimilar to known nuclear localization signals. This suggests that there might be other factors involved in its nuclear transport machinery (Nakayama et al., 2005). Moreover, we described a novel phosphorylation site (S635) in murine SATB1 protein. The S635A mutation resulted in SATB1’s aggregation and abruption of its DNA binding ability. Thus, it is reasonable to hypothesize that phosphorylation is also crucial for SATB1’s interactions with RNA. SATB1 is the substrate for a great number of post-translational modifications (Zelenka and Spilianakis, 2020), hence they likely serve important roles in the regulation of SATB1-dependent physiological processes.

The predicted saturation concentration (i.e. the concentration threshold that has to be crossed to trigger phase separation) of human SATB1 is 114 µM, which falls at the lower end of identified proteins with LLPS potential (Wang et al., 2018). However, the low saturation concentration would allow a precise regulation of SATB1’s phase transitions just by modulating its expression levels. Moreover, given the different biophysical properties of the two SATB1 isoforms, controlled production of either the short or the long SATB1 isoform would establish yet another layer of regulation, permitting tight regulation of SATB1’s biophysical behavior. We have recently described the SATB1-dependent promoter-enhancer network which controls the expression of important master regulator genes in developing T cells (Zelenka et al., 2022). Here we hypothesize, that the increasing protein levels of SATB1 and/or the long-to-short isoform ratio could promote the incorporation of the regulated genes into the transcriptional condensates, as it was also described for other factors (Cai et al., 2019; Zamudio et al., 2019; Lu et al., 2020). Such a mechanism of action would only be possible in primary cell types with high SATB1 concentration, such as in DP T cells, in embryonic stem cells and/or in some neurons.

The existence of two SATB1 protein isoforms with distinct biophysical properties has a clear physiological importance. However, the presence of the long isoform may also determine the pathological outcome in a number of disorders. Moreover, deregulation of any of the described mechanisms, such as the altered RNA binding capacity of SATB1, deregulation of its multivalency by mutations or by post-translational modifications, deregulation of its expression levels, or changes to the epigenetic landscape, ultimately leading to a different proportion of the long and short SATB1 isoforms, could all result in physiological aberrations. Future studies should draw their attention to the isoform-specific expression levels of SATB1, ideally coupled with microscopy, to reveal its localization patterns. This is particularly important given the higher phase separation propensity of the human long SATB1 isoform compared to the murine SATB1 (**Figure 5d**). Therefore, human cells could be more susceptible to the formation of aggregated SATB1 structures which could be associated with physiological defects. In line with our findings, nuclear SATB1 was associated with favorable outcome of cancer patients, whereas cytoplasmic SATB1 with poor prognosis (Durślewicz et al., 2021). Moreover, more focus should be brought to potential mutations in the susceptible poly-Q region of SATB1. Mutations of the poly-Q region may be difficult to identify, however they could lead to the extension of the poly-Q stretch, ultimately facilitating aggregate formation and thus a pathological situation as known for other PrLD-bearing proteins (Gruber et al., 2018; Peskett et al., 2018).

## 5 Data Availability Statement

RNA-seq experiments and SATB1 binding sites are deposited in Gene Expression Omnibus database under accession number GSE173470 and GSE173446, respectively.

## 6 Author contributions

T.Z. and C.S. designed the study. T.Z. cloned the CRY2 constructs and transfected the cells, performed the microscopy experiments, RNA-seq experiments and the computational analyses. D.A.P. performed the immunodepletion, western blot and LLPS centrifugation experiments. P.T. created the *Satb1* cKO mouse, cloned the original *Satb1* transcripts and generated custom-made antibodies. G.P. performed the SATB1 co-immunoprecipitation and mass spectrometry experiments. K.C.T. analyzed the mass spectrometry data. V.M.P. and T.Z. performed and analyzed the Raman spectroscopy experiments. G.Pap. performed immunofluorescence experiments. E.M. SATB1 protein production. M.K. Turbidity assay. J.P. performed FRAP experiments. D.S. provided resources for the super-resolution microscopy and consulted the imaging experiments and the analysis for alternative splicing. T.Z. wrote the original manuscript. C.S. supervised the work, obtained funding and corrected the manuscript. All authors read, discussed and approved the manuscript.

## 7 Funding

This work was supported by H2020-MSCA-ETN-2014 (GA642934) (TZ, CS), FONDATION SANTE X-COAT (TZ, CS), Ministry of Education Youth and Sports of the Czech Republic (LM2018129, CZ.02.1.01/0.0/0.0/18_046/0016045) (68378050-KAV-NPUI) (LTAUSA18103) (DS). The funders had no role in study design, data collection and analysis, decision to publish, or preparation of the manuscript.

## 8 Acknowledgements

We would like to thank Despina Tsoukatou for fruitful discussions and assistance in the lab. This work was supported by the European Union (European Social Fund ESF) and Greek national funds through the Operational Program ‘Education and Lifelong Learning’ of the National Strategic Reference Framework (NSRF) Research Funding Program ARISTEIA [MIRACLE 42], by FONDATION SANTE (X-COAT) and by Chromatin3D-H2020-MSCA-ITN (GA642934). Author V.M.P acknowledges the support of Stavros Niarchos Foundation within the framework of the project ARCHERS (“Advancing Young Researchers’ Human Capital in Cutting Edge Technologies in the field of Systems Biology Approaches and Personal Genomics for Health and Disease Treatment”). We acknowledge the Light Microscopy Core Facility, IMG CAS, Prague, Czech Republic, supported by MEYS (LM2018129, CZ.02.1.01/0.0/0.0/18_046/0016045) and RVO: 68378050-KAV-NPUI, for their support with the super-resolution imaging and image analysis presented herein. We thank the personnel of the Proteomics Facility at the Institute of Molecular Biology and Biotechnology, M. Aivaliotis and N. Kountourakis, for the Mass spectrometry analysis of tryptic peptides by nanoflow liquid chromatography with tandem mass spectrometry. Moreover, the results shown here are in part based upon data generated by the TCGA Research Network: https://www.cancer.gov/tcga. The funders had no role in study design, data collection and analysis, decision to publish, or preparation of the manuscript.

## 9 Conflict of interest

The authors declare that the research was conducted in the absence of any commercial or financial relationships that could be construed as a potential conflict of interest.

## Notes

### Competing Interest Statement

The authors have declared no competing interest.

### Summary of Updates

New data added; manuscript re-written; author affiliations updated; Supplemental files updated.

