## Supplementary Material for "A novel SATB1 protein isoform with different biophysical properties"

#### 1 Supplementary Methods

##### 1.1 2D gel electrophoresis coupled to Western Blot

Protein extracts from murine thymocytes (250 µg) were mixed with rehydration buffer (7 M Urea, 2 M Thiourea, 1% NP-40, 1% IPG ampholytes, 40 mM DTT, 0.25 mg bromophenol blue) in a final volume of 200 µl. The mix was spread throughout the length of a lane of the focusing tray and an 11 cm IPG strip (pH range 5.5-6.7) was slowly deposited upon the sample with the protein extract. The IPG strip was actively rehydrated with the application of 50 V for 16 hours. After the rehydration, wet filter wicks were added below the IPG strip upon the electrodes and the IPG strips were focused according to the manufacturer's suggestions for the specific IPG strip (Step 1: 250 V for 15 minutes in Rapid Ramp, Step 2: 8000 V for 1 hour in Slow Ramp, Step 3: 8000 V and 35,000 V-hr in Rapid Ramp). Upon isoelectric focusing the IPG strips were equilibrated for 15 minutes in Equilibration Buffer I (6 M Urea, 2% SDS, 0.375 M Tris pH 8.8, 20% Glycerol, 130 mM DTT) and then in Equilibration Buffer II (6 M Urea, 2% SDS, 0.375 M Tris pH 8.8, 20% Glycerol, 135 mM iodoacetamide). Strip was rinsed in 1× Running Buffer (25 mM Tris, 192 mM Glycine, 0.1% SDS) and loaded on top of a vertical 10% SDS-polyacrylamide gel. The well was sealed using agarose sealing solution (0.5% agarose, 0.002% bromophenol blue, 1× Running Buffer) and the gel was run at 230 V for 2 hours. The gel was transferred in a nitrocellulose membrane for 1.5 hours at 340 mA. The membrane was blocked using 5% nonfat milk/TBS-T and blotted with an antibody against SATB1 in a dilution 1:200 (Santa Cruz Biotechnology, sc-376097) and a donkey anti rabbit-HRP (Jackson Laboratories, dilution 1:2500).

##### 1.2 Recombinant SATB1 isoforms expression and purification

For SATB1 isoforms expression, the coding sequences of each isoform was cloned in a pRSETa bacterial expression vector that enabled the expression of the desired proteins fused with a 6×His tag. For large scale protein preparation, transformed BL21(DE3) pLysS competent cells were induced for SATB1 isoform expression with 0.25 mM IPTG for 16 hours at 18°C. Cell pellets were resuspended in 10-fold the pellet weight in Urea Lysis Buffer (8 M Urea, 10 mM Tris pH 8.0, 100 mM NaxPO4 pH 7.0, 10 mM Imidazole, 10 mM β-mercaptoethanol) and sonicated for 3 minutes at 15 Watts on ice using a probe sonicator. Samples were centrifuged for 20 minutes at 14,000 rpm and the supernatant was mixed with Ni-NTA agarose beads and incubated at room temperature for 3-6 hours with rotation. After the binding of the his-tagged proteins, beads were washed three times with Urea Lysis Buffer and eluted for 15 minutes with consecutive incubation in Urea Lysis Buffer (8 M Urea, 10 mM Tris pH 8.0, 100 mM NaxPO4 pH 7.0, 10mM Imidazole, 10 mM β-mercaptoethanol) that contained increasing amounts of Imidazole (100 mM, 250 mM, 0.5 M, 1 M and a final elution with 10 mM EDTA). Protein purification was verified in a denaturing polyacrylamide gel using Coomassie Brilliant Blue stain and Western blotting using antibodies that specifically recognize either the long isoform or both the long and short isoform. Purified proteins were dialyzed in dialysis tubing with consecutive dilution of the Urea concentration from 8-0 M in Dialysis Buffer (20 mM Tris pH 8.0, 10% glycerol, 300 mM NaCl, 1 mM DTT, 0.1% NP40).

##### 1.3 Quantification of Phase Separation (Sanulli and Narlikar, 2021)

Centrifugation assay: Purified long and short SATB1 isoforms were diluted in Phase Separation Buffer (20 mM HEPES pH 7.8, 150 mM NaCl, 1 mM DTT) and mixed with equal volume of 1.5 μg RHS6 DNA or empty vector pBluescript DNA. Samples were gently mixed, incubated at room temperature for 20 minutes and centrifuged for 10 minutes at 10,000 g at room temperature. The amount of DNA left in the supernatant was measured in triplicates at 260 nm.
Turbidity assay: In 1.5-ml tubes containing 15 μl each of the different SATB1 stock solution (8, 4, 3, or 2 μM) were added 15 μl of DNA (RHS6 or control pBluescript 2× stock solution, 100 ng/μl in 10 mM Tris-HCl pH 8, 0.5 mM EDTA pH 8). In 1.5-ml tube, 15 μl of DNA 2× stock solution was supplemented with 15 μl SATB1 buffer (20 mM HEPES pH 7.8, 150 mM KCl, 1 mM DTT). The latter was the reference for 100% solubility (no phase separation), as DNA is in solution under the described conditions in the absence of SATB1. Four more 1.5-ml tubes containing 15 μl each of the different SATB1 stock solution (8, 4, 3, or 2 μM) were supplemented with 15 μl of TE0.5 (10 mM Tris-HCl pH 8, 0.5 mM EDTA pH 8) instead of DNA to each tube. These samples were used to verify the contribution of SATB1 to the 260 nm absorbance. The solution was mixed gently and incubated for 20 minutes at room temperature. 20 μl of each sample were transferred to a clear-bottom 384-well plate. 20 μl phase separation buffer was added to a separate well, to use as negative control. The plate was sealed with a film to prevent sample evaporation and absorbance was measured at 340 nm in a plate reader.

##### 1.4 RT-qPCR

Isolated thymocytes were resuspended in 1 ml of TRIzol Reagent (Sigma Aldrich, T9424) and RNA was isolated according to the manufacturer's protocol. Isolated RNA (5 μg) was decontaminated by incubation with 4 units of DNase I (NEB, M0303L) at 37°C for 30 minutes. The reaction was stopped by adding 10 μl of DNase Inactivation Reagent (Ambion, 8174G) and incubating at 70°C for 10 minutes. The supernatant was transferred to a new tube and RNA was precipitated by adding 10 μg of Linear Acrylamide (Ambion, AM9520), 2.5X volumes of 100% Ethanol and incubated at –80°C for 30 minutes. The supernatant was removed and the pellet was washed twice with 75% Ethanol. The air-dried pellets were resuspended in RNase-free water and incubated at 55°C for 15 minutes to dissolve. For reverse transcription, 1 μg of RNA was combined with 1 μl of oligo dTs (100 pmol/μl), 1 μl dNTPs (10 mM) and topped up with $ddH_2O$ to 13 μl. Samples were denatured at 65°C for 5 minutes and transferred on ice. Samples were combined with 4 μl 5X Minotech RT assay buffer, 1 μl 0.1 M DTT, 1 μl RNase inhibitor (40 U/μl) and 1 μl Minotech RT enzyme (Minotech, 801-1). The mixture was incubated at 45°C for 60 minutes and then heat-inactivated at 70°C for 15 minutes. The original transcripts were quantified with qPCR, using SYBR Select Master Mix (Applied Biosystems, 4472908) according to the manufacturer's instructions with the following primers: long *Satb1* isoform (fwd: TTCTTACCAAGCCTGCTGACC, rev: TGGTCTTCTGCCGGTTTTCC), all short *Satb1* isoforms (fwd: CAGAACAGATTCAGCAGCAG, rev: TGGTCTTCTGCCGGTTTTCC) and *Dnmt1* (fwd: GCAGCCTATGAAGATTGAAGAG, rev: CCTGGCTAGATAGAGTGGTTTG). Three technical replicates from a male and female mouse were performed.

##### 1.5 Identification of SATB1's phosphosite S635

DNA affinity chromatography coupled with mass spectrometry was performed as previously described (Yaneva and Tempst, 2003, 2006) to identify associated proteins. A region of 151 bp, part of the RHS6 DNase I hypersensitive site of the TH2-LCR (Spilianakis et al., 2005; Williams et al., 2013) was cloned into TOPO-TA vector using following primers: RP1-fwd CTTCCATTTGCTGGCCTCT; RP2-rev CCTGCATCTGATACTTAGACGTA. Two types of biotinylated oligonucleotides were utilized. Targeted fragment was biotinylated by PCR amplification with the following primers: biotinT7-fwd Biotin-d(CTCGAGTAATACGACTCACTATAGG); M13-rev AACAGCTATGACCATG. The same primers were used to prepare oligonucleotide for the negative selection (preclearing), by amplification from the TOPO-TA vector without the RHS6 region. 1 mg of M-280 Streptavidin Dynabeads (Invitrogen, 11205D) were resuspended and washed three times according to the manufacturer's instructions. 100 μl of 2× Binding & Wash buffer (10 mM Tris pH=7.5, 1 mM EDTA, 2 M NaCl) containing 10 μg of the biotinylated oligonucleotide in an equal volume of water was added. The beads with the oligonucleotide were incubated for 15 minutes with rotation at room temperature. Beads were collected on a magnet and further washed with 1× Binding & Wash buffer and once with DNA binding buffer (20 mM HEPES pH=7.9, 0.1 mM KCl, 0.2 mM EDTA, 0.01% NP-40, 0.5 mM DTT, 0.2 mM PMSF, 10% Glycerol) supplemented with 0.1 mg/ml oligo dI:dC and poly dI:dC. Thymocyte nuclear extracts (5 mg) were first negatively selected by the "preclearing" biotinylated oligonucleotide and then positively selected utilizing the biotinylated RHS6 sequence. The biotinylated oligonucleotides used for the positive selection of bound thymocyte proteins were boiled and electrophoresed on a denaturing polyacrylamide gel. Upon Coomassie blue silver staining, individual slices of the positively selected samples, the negatively selected samples and washes were utilized in parallel for nanoLC-MS/MS analysis as previously described (Stratigi et al., 2015). Subsequent protein identification was done using Mascot and Sequest software.

##### 1.6 Raman spectroscopy

Isolated thymocytes were washed once with 1× PBS and then incubated in 10% 1,6-hexanediol (Sigma Aldrich, 240117) in 1× PBS solution (Sigma Aldrich, P4417) for 10 minutes at room temperature, side by side with a non-treated control. Samples were fixed with 1% formaldehyde (final conc.; Pierce, 28908) for 10 minutes and then quenched with glycine to a 0.125 M final concentration and incubated at room temperature for 5 minutes, while rocking. Cells were washed five times with 1× PBS. For measurements, a drop of cells was loaded on a stainless-steel microscopic slide (Renishaw, UK) and imaged through an Olympus 60x water immersion objective lens with a numerical aperture (NA) of 1.2 and a working distance of 280 μm (UPLSAPO60XW/1.2, Olympus, Tokyo, Japan). Raman measurements were made by a modified Raman Spectrometer (LabRAM HR, HORIBA Scientific, Lille, France). The excitation laser line used had a central wavelength at 532 nm and the resulting laser power on the sample was ~10 mW. The laser spot spatial diameter was approximately 0.6 μm, with an axial length of about 0.5 μm. A grating of 600 groves was used that resulted in a Raman spectral resolution of around 2 $cm^{-1}$. A temperature-controlled stage (PE120-XY, Linkam) was coupled with the microscope stage that kept cells temperature constant. Spectral calibration was performed with a $SiO_2$ reference sample, presenting a single peak at 520.7 $cm^{-1}$. Raman signal was acquired in the spectral range between 300 and 3,150 $cm^{-1}$. Acquisition time was set to 10 seconds with a spectral accumulation of 3 spectra/point. All Raman spectra acquired followed this processing methodology: a) cosmic rays were removed by an internal function (Despike, LabSpec LS6, Horriba); b) background signal was subtracted from the raw Raman spectral data using a polynomial function; c) all spectra were normalized in intensity by an internal function (Unit Vector, LabSpec LS6, Horriba). PCA was performed using the Raman PCA Tool (Candeloro et al., 2013), publicly available through (https://sourceforge.net/projects/ramantoolset/). For each condition, 2-5 biological replicates were used.

#### 1.7 Tissue cultures and transient transfection

Cells were cultured in the complete growth medium DMEM (Gibco, 11995-073) supplemented with 10% Fetal Bovine Serum (Sigma Aldrich, F4135) and with 2 mM L-glutamine (Gibco, 25030) in a 5% $CO_2$ incubator at 37°C, until 70% confluency. Before the experiment, cells were washed in Dulbecco's PBS (DPBS; Gibco, 14200067) without $Mg^{2+}$ and $Ca^{2+}$. Cells were treated for 3-5 minutes at 37°C in 1× trypsin solution until detached. Trypsin was quenched by complete growth medium and cells collected. For optoDroplet experiments, 800 ng of plasmid DNA was combined with 50 µl of Opti-MEM I Reduced Serum Medium and pipetted onto a glass bottom dish (MatTek, P35GC-0-14-C). Next, 2 µg of Lipofectamine 2000 (Invitrogen, 11668-027) was carefully mixed with 50 µl Opti-MEM I Reduced Serum Medium and it was incubated for 5 minutes at room temperature. Lipofectamine/OptiMEM mix was slowly and drop-wise added to the DNA/OptiMEM mixture on the glass bottom dish and incubated for 30 minutes at room temperature. 50,000 of NIH/3T3 ATCC CRL-1658 cells were added to the DNA/Lipofectamine/OptiMEM mixture and they were incubated at 37°C in a humidified 5% $CO_2$ incubator for 6 hours. After six hours, 2 ml of the growth medium with antibiotics (penicillin/streptomycin to the final concentration 100 I.U./ml / 100 µg/ml) was added and continued transfection until ready to assay (24-48 h post transfection).

### 2 Supplementary Figures

#### Supplementary Figure 1

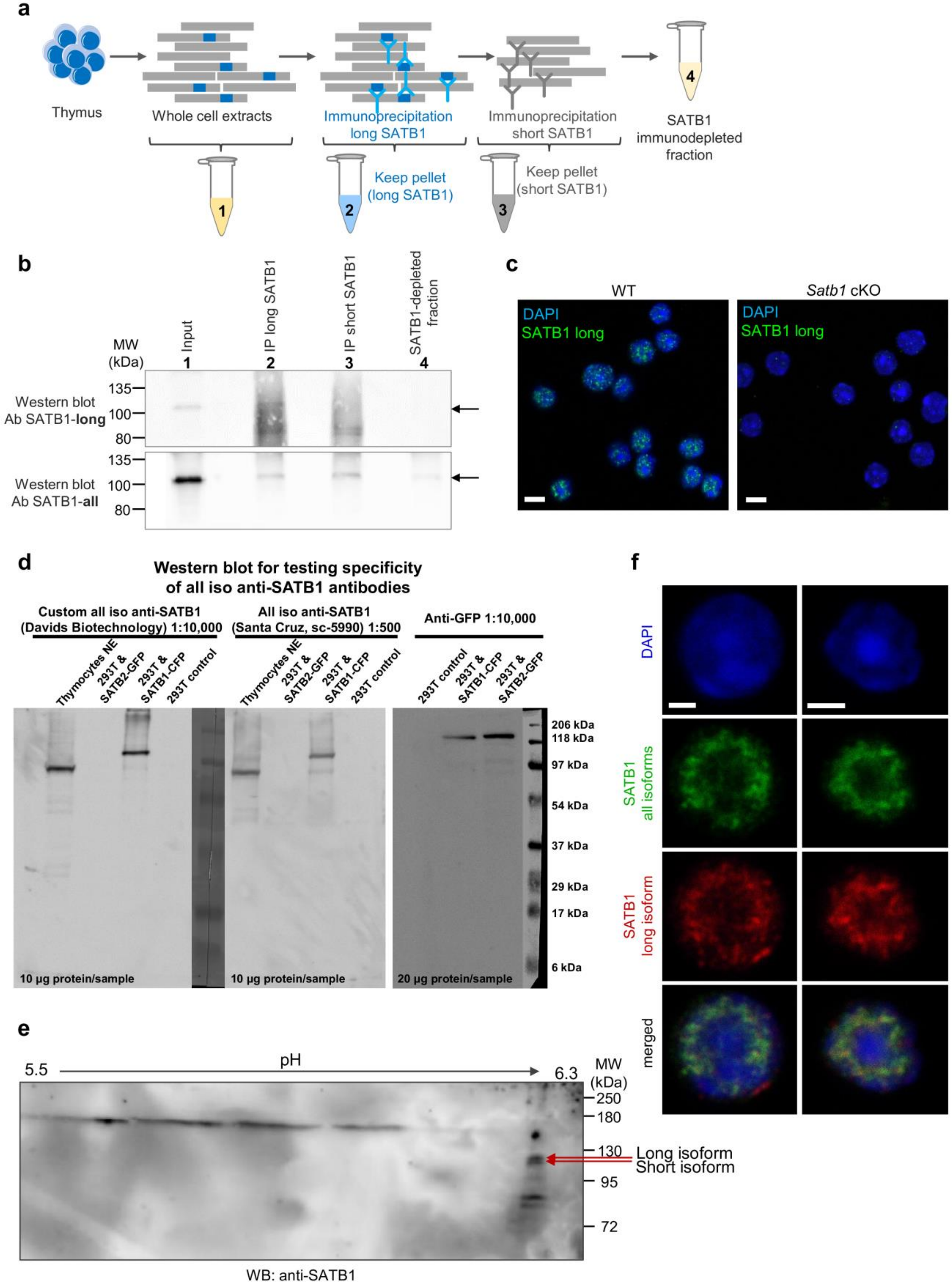

**Supplementary Figure 1. Validation of SATB1 antibodies and long SATB1 isoform existence. a**, Scheme for the approach utilized to detect SATB1 long isoform and validate the custom-made long isoform-specific antibody (Davids Biotechnology). Whole cell thymocyte protein extracts (sample 1) were prepared and incubated with a custom-made antibody against the long SATB1 isoform. The immunoprecipitated material (sample 2) was kept and the immunodepleted material was subjected on a second immunoprecipitation reaction utilizing an antibody detecting epitopes on both long and short SATB1 isoforms (Santa Cruz Biotechnology, sc-376096). The material from the second immunoprecipitation reaction was kept (sample 3), as well as the immunodepleted thymocyte extract for all SATB1 isoforms (sample 4). **b**, Western blot analysis for the samples described in **a**. The whole thymocyte protein extract (1), immunoprecipitated long SATB1 protein (2), immunoprecipitated short SATB1 protein (3) and total thymocyte protein fraction immunodepleted for both SATB1 isoforms (4). UPPER PANEL: Western blot analysis utilizing a SATB1 antibody detecting only the long SATB1 isoform. LOWER PANEL: Western blot analysis utilizing a SATB1 antibody detecting all SATB1 isoforms. **c**, Representative single z-stack images of immunofluorescence experiments performed with primary murine thymocytes and analyzed with confocal microscopy utilizing the custom-made polyclonal antibody raised against the extra SATB1 peptide of the long isoform. The long SATB1 isoform is detected (green) in C57BL/6 thymocytes but not in *Satb1* cKO thymocytes. Scale bar 5 μm. **d**, Western blot analysis comparing the specificity of the custom-made antibody targeting all SATB1 isoforms (Davids Biotechnology) and the commercial antibody (Santa Cruz Biotechnology, sc-5990), both targeting the N-terminus of SATB1. Murine thymocyte nuclear extracts were compared with extracts from transfected 293T cell line. Both antibodies showed high specificity towards SATB1. **e**, 2D gel electrophoresis coupled to western blotting for SATB1. Isoelectric focusing of murine thymocyte protein extracts, run in SDS PAGE and transferred to nitrocellulose blotted with an anti-SATB1 antibody detecting the long and short SATB1 isoforms. **f**, Representative single z-stack images of immunofluorescence experiments analyzed with confocal microscopy utilizing a commercially available antibody against SATB1 detecting both short and long isoforms (C-6: sc-376096, mouse monoclonal IgG1κ, Santa Cruz Biotechnology) and the custom-made polyclonal antibody raised against the extra SATB1 peptide of the long isoform. Scale bar 2 μm.

#### Supplementary Figure 2

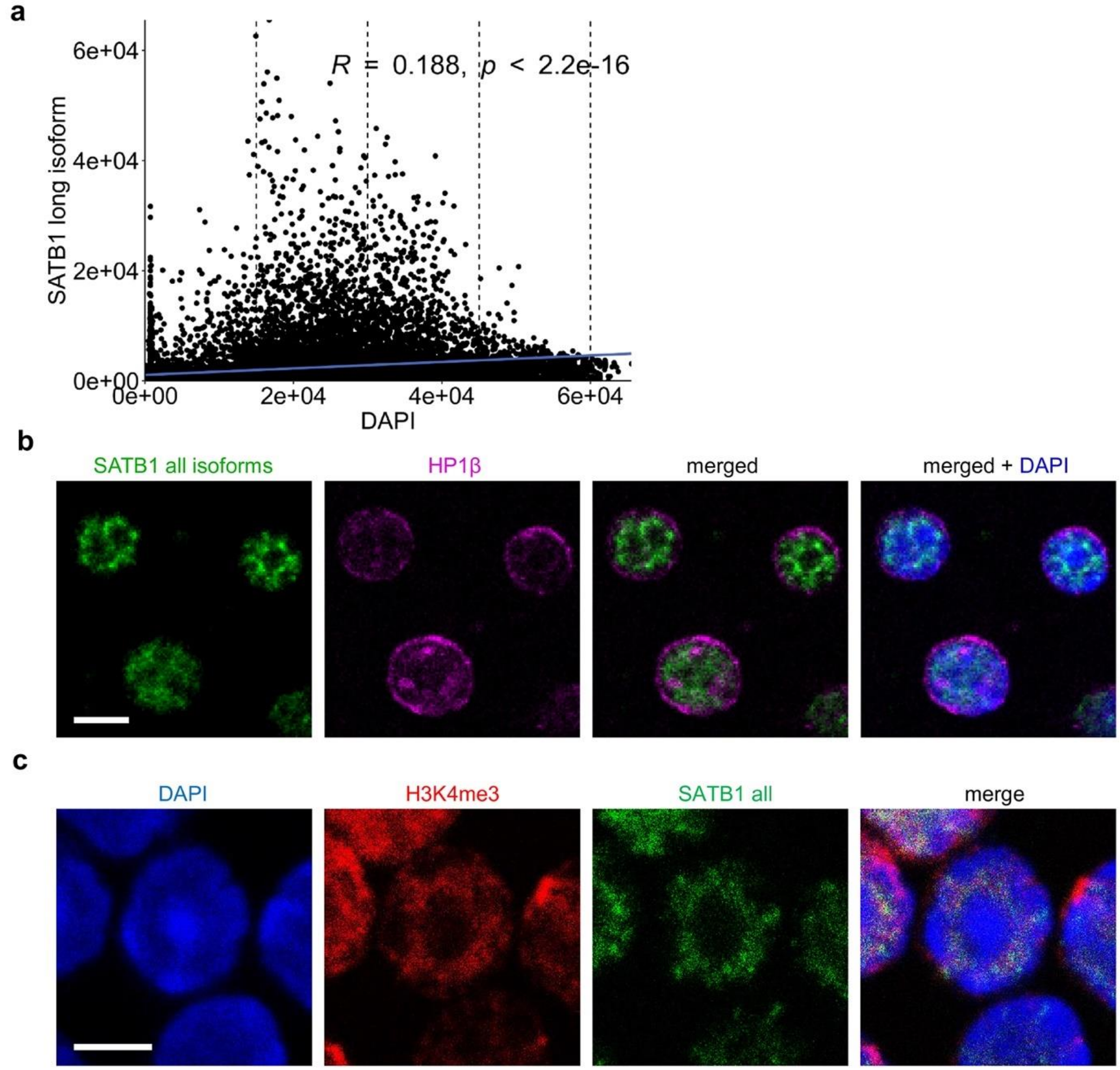


**Supplementary Figure 2. SATB1 is positively correlated with nuclear regions with transcriptional activity. a**, Fluorocytogram of the pixel-based co-localization analysis (based on 3D-SIM) between the long SATB1 isoform and DAPI signal, indicating the localization of SATB1 in different nuclear zones. The three vertical dashed lines categorize the DAPI signal to highlight the different proportions of SATB1 signal in different nuclear zones. **b**, SATB1 shows a negative correlation with heterochromatin mark HP1β. Representative single z-stack images of immunofluorescence experiments performed with primary murine thymocytes and analyzed with confocal microscopy utilizing a commercially available antibody against SATB1 (Santa Cruz Biotechnology , sc-5990, dilution 1:50, secondary donkey anti goat Alexa 647, dilution 1:250) and anti-HP1β (Santa Cruz Biotechnology , sc-20699, dilution 1:50, secondary donkey anti rabbit Alexa 546, dilution 1:250). Scale bar 5 µm. **c**, SATB1 shows a positive correlation with H3K4me3 histone mark of actively transcribed chromatin. The same experiment as in **b** but utilizing a commercially available antibody against SATB1 detecting both short and long isoforms (Santa Cruz Biotechnology, C-6: sc-376096, mouse monoclonal IgG1κ, dilution 1:100; secondary Alexa 488 goat anti mouse in a dilution of 1:250) and anti-H3K4me3 (Abcam - ab8580, dilution 1:500, secondary Alexa-647 goat anti rabbit, dilution 1:250). Scale bar 2 μm.

**Supplementary Figure 3**

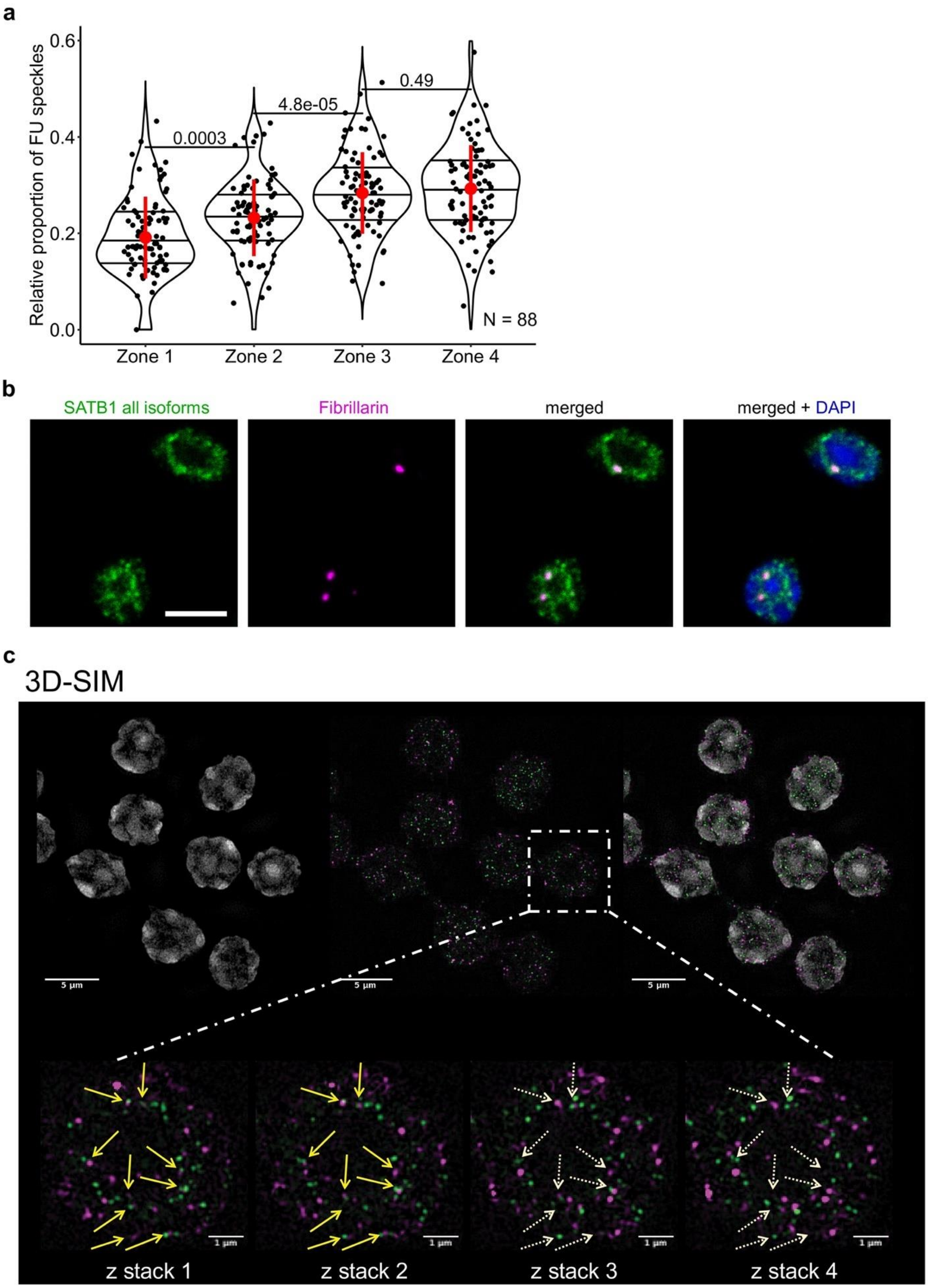

**Supplementary Figure 3. SATB1 associates with sites of active transcription. a**, Quantification of the 3D-SIM immunofluorescence images, related to Fig. 2b. Nuclei of primary murine thymocytes were categorized into four zones based on the intensity of DAPI staining and FU speckles in each zone were counted. Images used represented a middle z-stack from the 3D-SIM experiments. The horizontal lines inside violin plots represent the 25$^{th}$, 50$^{th}$ and 75$^{th}$ percentiles. Red circles represent the mean ± s.d. *P* values by Wilcoxon rank sum test. **b**, SATB1 co-localizes with a nucleolar protein Fibrillarin. Representative single z-stack images of immunofluorescence experiments performed with primary murine thymocytes and analyzed with confocal microscopy utilizing a commercially available antibody against SATB1 (Santa Cruz Biotechnology, sc-5990, dilution 1:200, secondary donkey anti goat Alexa 647, dilution 1:250) and anti-Fibrillarin (Abcam, ab5821, dilution 1:300, secondary donkey anti rabbit Alexa 546, dilution 1:250). Scale bar 5 μm. **c**, 3D-SIM microscopy experiment showed co-localization between the SATB1 long isoform and sites of active transcription, labeled with fluorouridine (FU). Four consecutive z-stacks are shown to better indicate that the majority of SATB1 speckles overlaid with FU staining, even though it could not be always apparent in a selected single stack. Scale bar 5 μm in the top and 1 μm in the bottom. Scale bar 5 μm in the top and 1 μm in the bottom panel.

#### Supplementary Figure 4

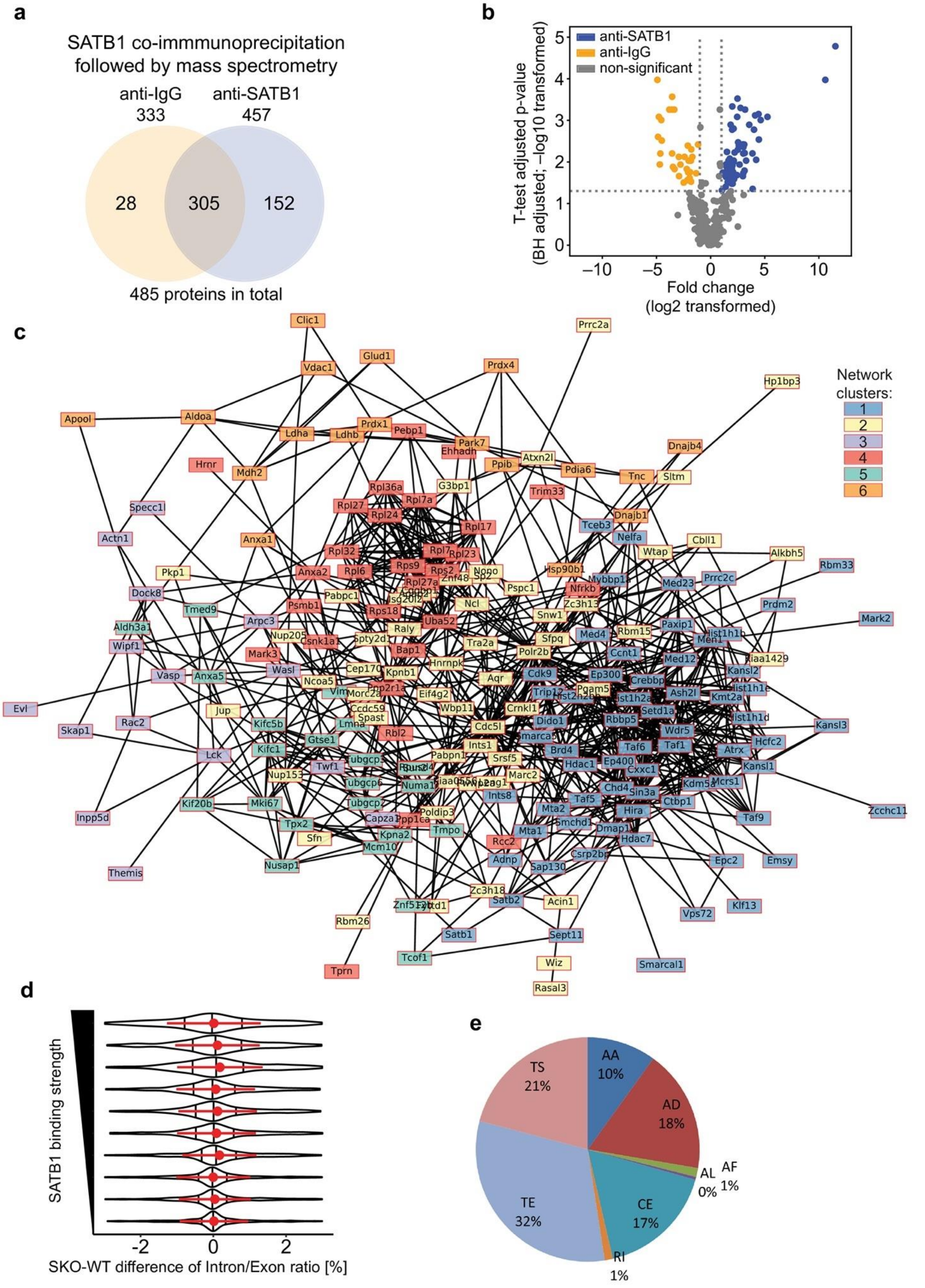

**Supplementary Figure 4. SATB1 interacts with proteins involved in transcription and splicing.** **a**, Overview of detected proteins in the SATB1 co-immunoprecipitation experiments followed by mass spectrometry analysis (IP-MS). IgG co-immunoprecipitation was used as a negative control. **b**, Volcano plot depicting the significance of proteins detected in the SATB1 IP-MS experiments. **c**, A full version of the k-means clustered STRING network of all significantly enriched SATB1-interacting proteins that was presented in Fig. 2d. **d**, Correlation between SATB1 binding and deregulation of splicing in *Satb1* cKO thymocytes, supporting the direct involvement of SATB1 in splicing. The vertical lines inside violin plots represent the 25$^{th}$, 50$^{th}$ and 75$^{th}$ percentiles. The red circles represent the mean ± s.d. **e**, Summary of groups for differential splicing events in *Satb1* cKO identified by Whippet (Sterne-Weiler et al., 2018) (for the analysis BAM files were utilized to detect unannotated splicing events). AA – Alternative Acceptor splice site, AD – Alternative Donor splice site, AF – Alternative First exon, AL – Alternative Last exon, CE – Core exon which may be bound by one or more alternative AA/AD nodes, RI – Retained intron, TE – Tandem alternative polyadenylation site, TS – Tandem transcription start site.

**Supplementary Figure 5**

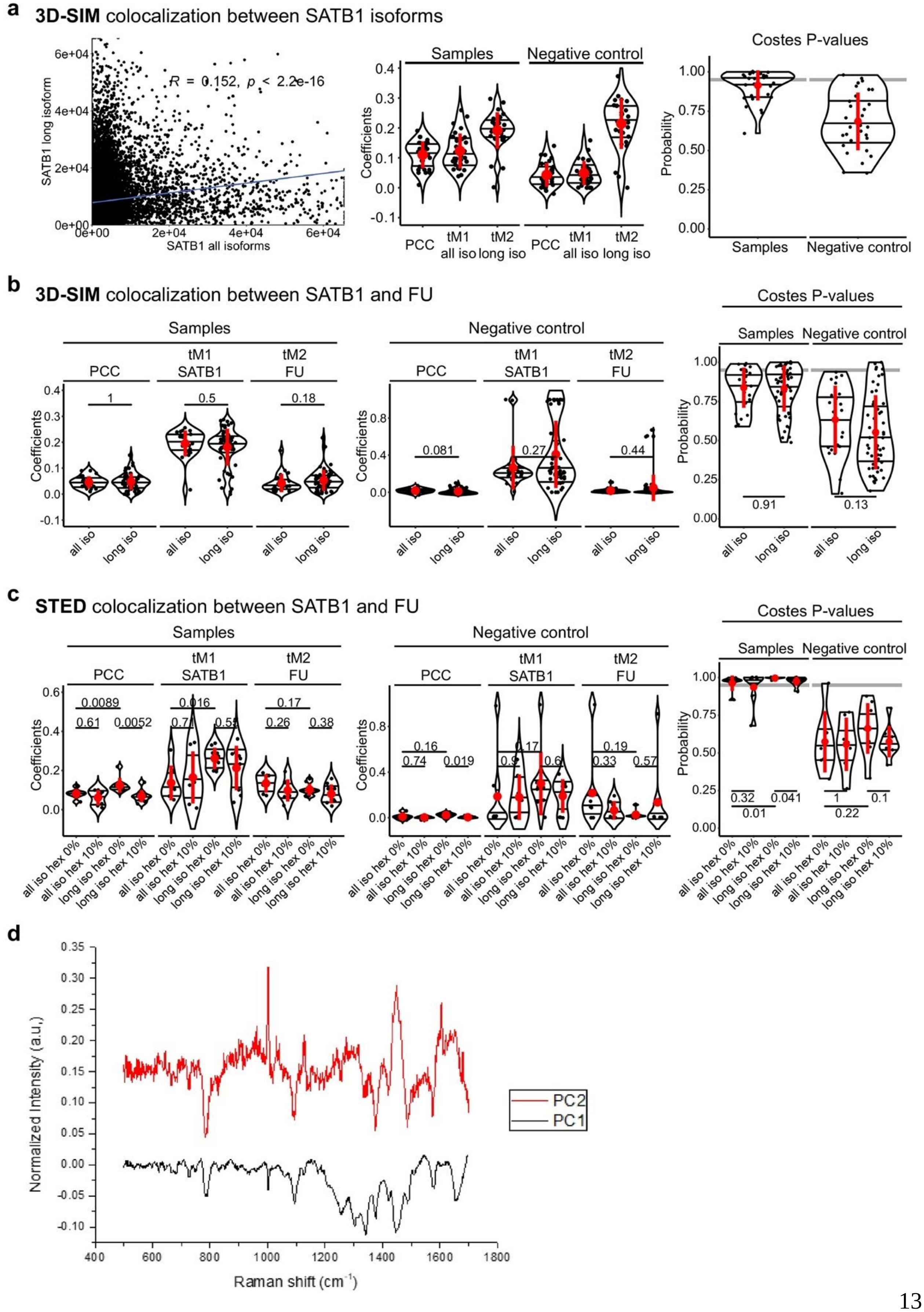

**Supplementary Figure 5. Analysis of pixel-based co-localization and Raman spectroscopy. a**, Pixel-based co-localization (based on 3D-SIM) between SATB1 isoforms detected by the antibody targeting all the isoforms (Santa Cruz Biotechnology, sc-5990) and only the long isoform (Davids Biotechnology, custom-made). This figure serves as a positive control for **b** and **c**. LEFT: Fluorocytogram of the pixel-based co-localization based on 3D-SIM. MIDDLE: Pearson correlation coefficients (PCC) and thresholded Manders' coefficients (tM) derived from the pixel-based co-localization. The Manders' coefficients indicate a portion of a factor co-localizing with the other studied factor. In this figure, the tM1 values indicate a portion of SATB1 all isoforms' signal that co-localized with the long isoform-specific signal and tM2 showing a portion of the long isoform-specific signal being co-localized with all SATB1 isoforms. RIGHT: The Costes P-values (Costes et al., 2004) derived from randomly shuffled chunks of analyzed images (100 randomizations). Grey line indicates 0.95 level of significance. **b**, The same analysis as in **a** but applied on 3D-SIM images of SATB1 and fluorouridine (FU)-labeled sites of active transcription. **c**, The same analysis as in **a** but applied on STED images of SATB1 and FU-labeled sites of active transcription with and without 10% 1,6-hexanediol treatment. In **a**, **b** and **c**, the horizontal lines inside violin plots represent the $25^{th}$, $50^{th}$ and $75^{th}$ percentile. Red circles represent the mean ± s.d. *P* values by Wilcoxon rank sum test. **d**, Extracted Raman spectra for the two main principal components from the PCA analysis depicted in Fig. 3h.

#### Supplementary Figure 6

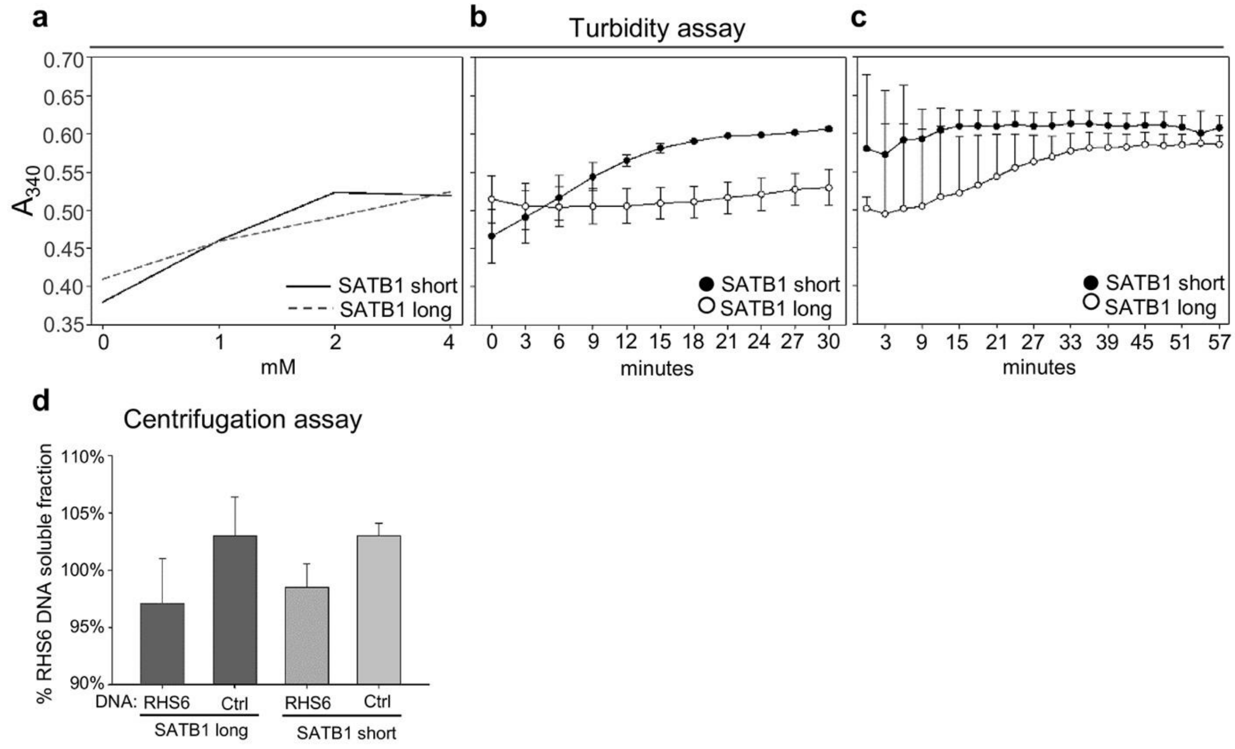


**Supplementary Figure 6. Quantitation of LLPS utilizing recombinant SATB1 isoforms proteins**. **a**, Turbidity of SATB1 assemblies was measured with increasing SATB1 concentration upon 5 minutes of LLPS induction. **b**, Turbidity of SATB1 assemblies was measured for 1 μM of SATB1 protein in a time-course upon LLPS induction. **c**, Turbidity of SATB1-RHS6 DNA assemblies was measured for 1 μM of SATB1 protein in a time-course upon LLPS induction. **d**, Centrifugation assay. The sedimentation of SATB1-DNA assemblies is measured in the presence of two types of either RHS6 or control (pBluescript) DNA. The y-axis displays the DNA in solution after centrifugation, as measured by absorbance at 260 nm. SATB1 isoform concentration 1 μM. Presented values are corrected for the absorbance at 280 nm.

#### Supplementary Figure 7

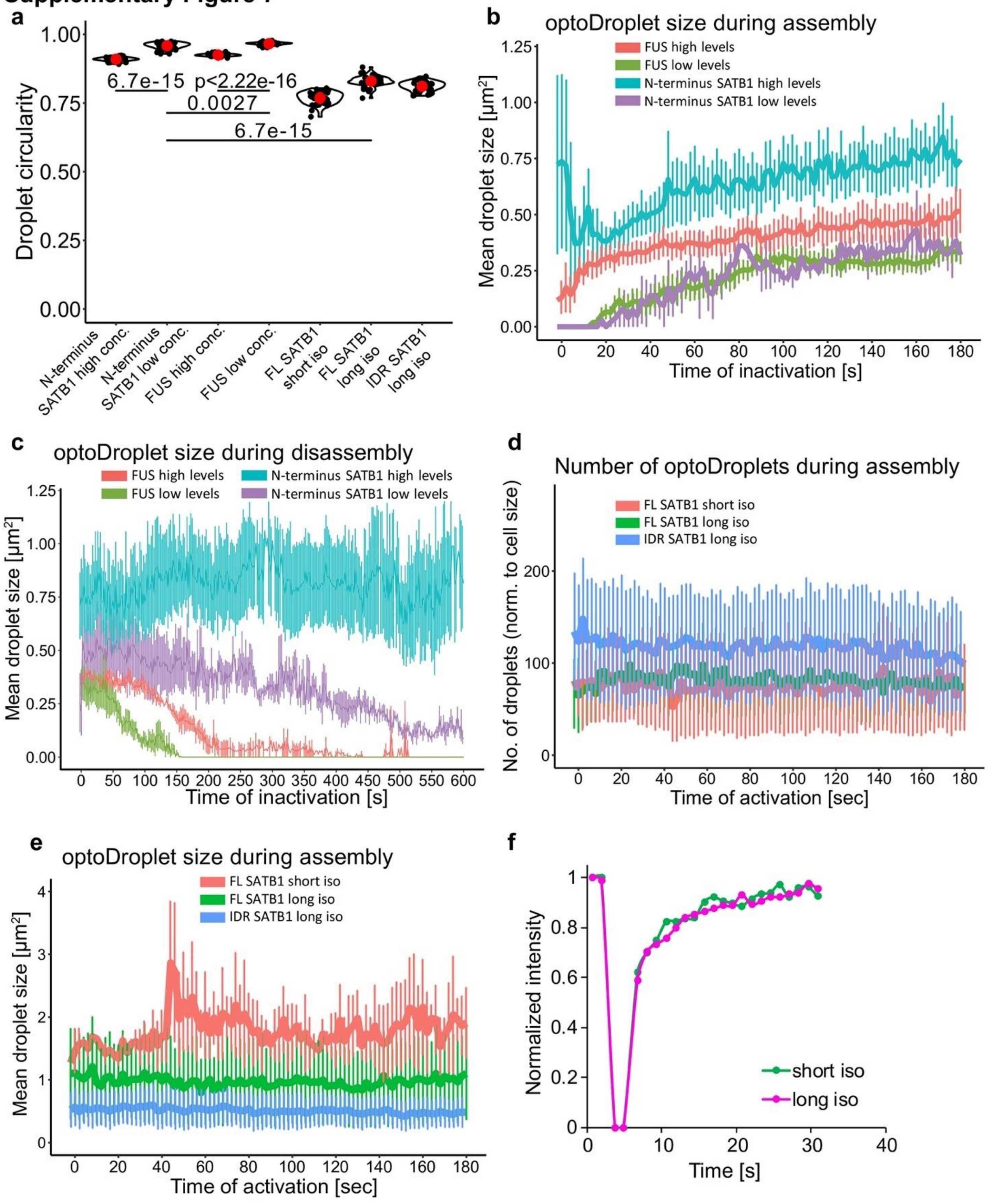


**Supplementary Figure 7. Analysis of optogenetics experiment results and full length SATB1 FRAP. a**, Droplet circularity for all analyzed optoDroplet constructs. Red circles represent the mean ± s.d. *P* values by Wilcoxon rank sum test. **b**, Average size of FUS and N-terminus SATB1 optoDroplets during activation measured as an area of the droplet in the analyzed z-stack. **c**, Average size of FUS and N-terminus SATB1 optoDroplets during the indicated inactivation time measured as an area of the droplet in the analyzed z-stack. **d**, Number of optoDroplets detected using the same approach as in Fig. 4d, utilizing full length (FL) and IDR SATB1 optoDroplet constructs. **e**, Average size of full length (FL) and IDR SATB1 optoDroplets during activation measured as an area of the droplet in the analyzed z-stack. In **b**, **c**, **d**, **e**, error bars represent the s.e.m. **f**, FRAP experiments on EGFP full length SATB1 short and long isoform constructs transfected into NIH-3T3 cells indicate similar molecular dynamics for the two isoforms in this *in vitro* system. The graphs show representative fluorescence recovery curves of the bleached areas in the nucleus over the shown time course (in seconds). The results are percentages of the postbleach over the prebleach intensities after correction for background and fluorescent decay during imaging. The error range was smaller than 20% of the mean values. The cells for live microscopy were grown in Lab-Tech chambers (Thermo Fisher Scientific), were transfected with the indicated plasmids and 48 h post-transfection were examined within 30 min at room temperature. Confocal microscopy was carried out on a Zeiss Axioscope 2 plus microscope equipped with a Bio-Rad Radiance 2100 laser scanning system and LaserSharp 2000 imaging software. Fluorescence recovery after photobleaching (FRAP) analysis was done using a standard region of interest (ROI) and monitoring close to saturation of recovery. After subtracting the background, fluorescent intensities were normalized against a companion unbleached ROI in the same cell.

#### Supplementary Figure 8

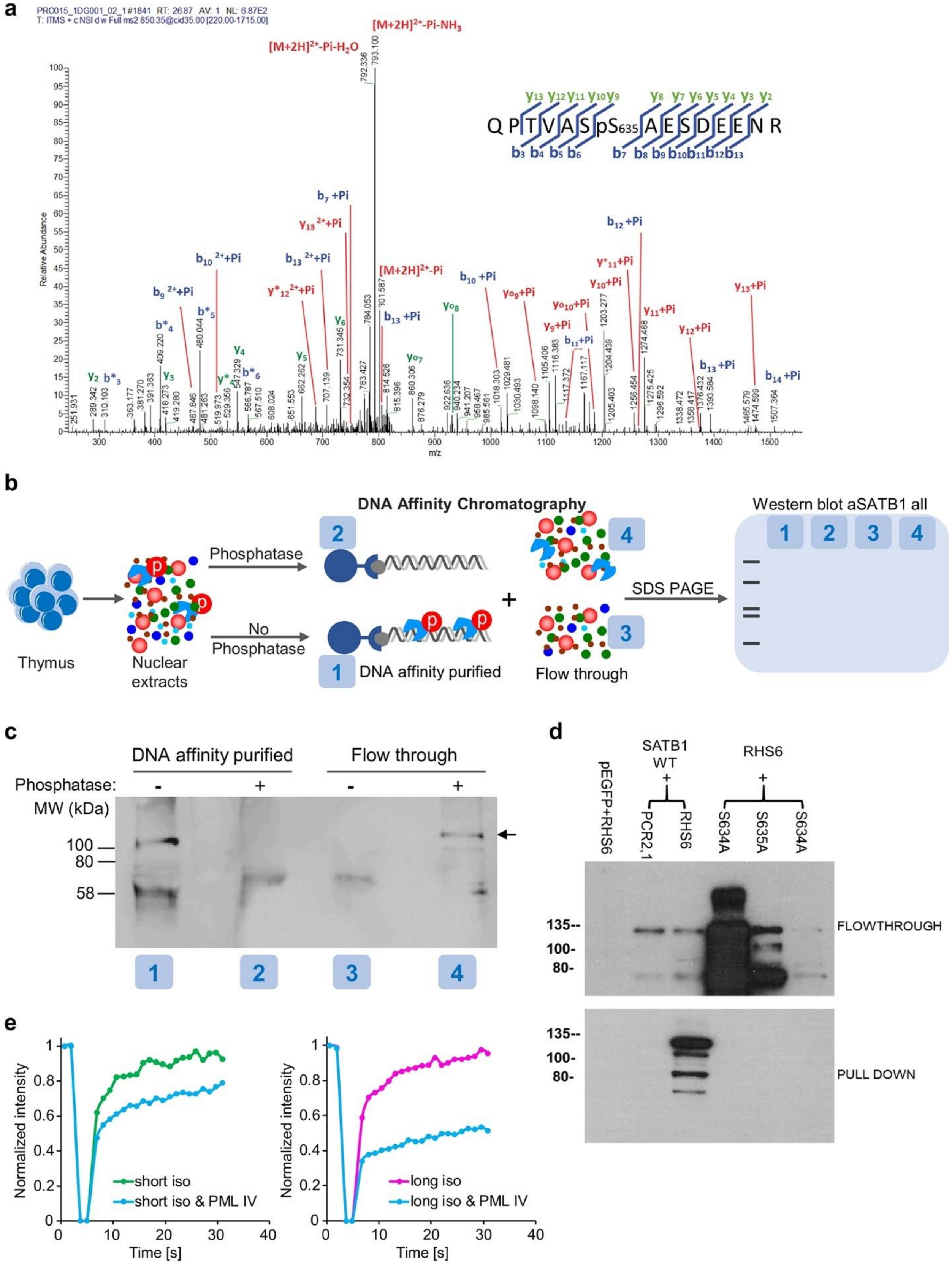

**Supplementary Figure 8. Identification and characterization of SATB1 novel phosphorylation site S635. a**, DNA affinity chromatography coupled to mass spectrometry using murine thymocyte protein extracts and an RHS6 biotinylated probe as a bait uncovered SATB1 S635 phosphosite. **b**, Scheme of DNA affinity purification experiment. Nuclear extracts from primary murine thymocytes were prepared and split in two fractions. One of the two samples was treated with phosphatase while phosphatase inhibitors were added in the other. DNA affinity purification followed, using biotinylated RHS6 DNA (RAD50 DNase I hypersensitive site of the TH2 Locus Control Region (Spilianakis et al., 2005; Williams et al., 2013), conserved between mouse and human) immobilized on streptavidin magnetic beads. The purified material was analysed with SDS-PAGE and blotted with Western blotting. **c**, Western blot results related to the scheme presented in **b**. The DNA bound material ("DNA affinity purified") and the non-bound ("Flow through") were analyzed with PAGE and blotted with a SATB1-specific antibody detecting all protein isoforms. The phosphatase-treated SATB1 protein did not bind RHS6 DNA. **d**, DNA affinity purification assay indicating that WT SATB1 effectively binds to the RHS6 site of the TH2 LCR gene (Spilianakis et al., 2005; Williams et al., 2013). In contrary, none of the SATB1 phosphomutants was pulled-down with the RHS6 DNA probe. The custom antibody targeting all SATB1 isoforms was used (Davids Biotechnology). **e**, FRAP experiments on EGFP full length SATB1 short and long isoform constructs transfected into NIH-3T3 cells as indicated in Supplementary Fig. 7f. In this experiment the constructs were co-transfected with PML IV isoform to demonstrate the impact of PML on SATB1 dynamics.

**Supplementary Figure 9**

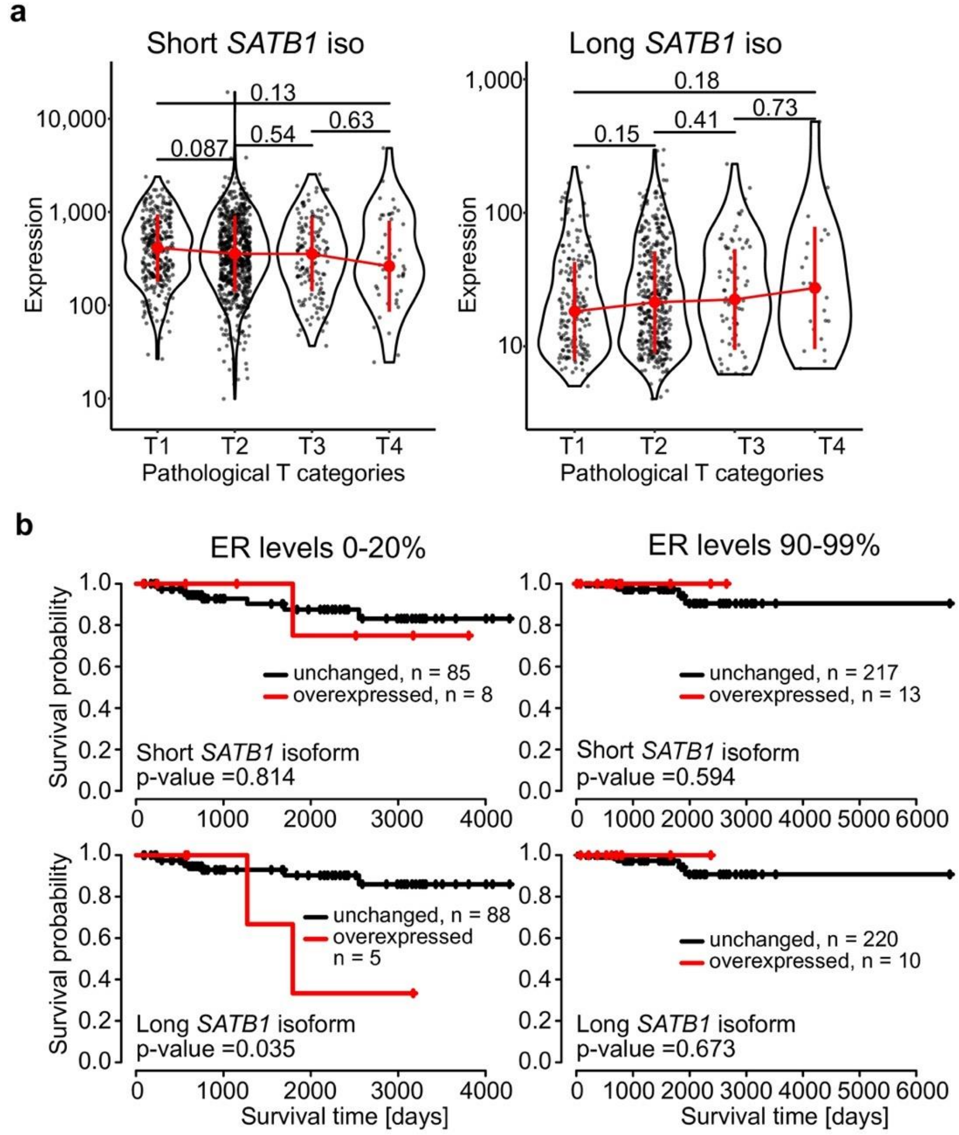


**Supplementary Figure 9. Long SATB1 isoform in human breast cancer patients is associated with worse prognosis and survival. a**, High pathological T categories of human breast cancer patients were associated with high expression of the long *SATB1* isoform. In contrast, the expression of the short *SATB1* isoform was negatively correlated with the pathological T categories. Red circles represent the median values ± s.d. *P* values by Wilcoxon rank sum test. **b**, Human breast cancer patients with low estrogen receptor levels and overexpression of the long *SATB1* isoform displayed worse survival. Overexpression of the short SATB1 isoform had a milder effect. In patients with high estrogen receptor levels, overexpression of any SATB1 isoform had a positive impact on survival. *P* values by chi-square test of the survival package in R.
